# Metabolite import via SLC33A1 enables ATF6 activation by endoplasmic reticulum stress

**DOI:** 10.1101/2025.07.17.664173

**Authors:** Ginto George, Heather P. Harding, Richard Kay, David Ron, Adriana Ordóñez

**Affiliations:** Cambridge Institute for Medical Research (CIMR), University of Cambridge, Biomedical Campus, The Keith Peters Building, Cambridge CB2 0XY, United Kingdom; Institute of Metabolic Science | Metabolic Research Laboratories, Addenbrooke’s Hospital. Hills Road, Cambridge, CB2 0QQ, United Kingdom; Universidad Católica de Murcia (UCAM), HiTech. Campus de los Jerónimos 135, E-30107, Guadalupe, Spain

**Keywords:** Acetylation, ATF6, CRISPR-Cas9 screen, Glycan sialylation, SLC33A1, UPR

## Abstract

The transcription factor ATF6 has a central role in adapting mammalian cells to endoplasmic reticulum (ER) stress via the Unfolded Protein Response (UPR). This has driven efforts to identify modulators of ATF6 signalling. Here, an unbiased genome-wide CRISPR-Cas9 screen performed in Chinese Hamster Ovary (CHO) cells revealed that proteolytic processing of the ATF6α precursor to its active form was impaired in CHO cells lacking the ER-resident solute carrier SLC33A1, a transporter involved in acetyl-CoA import, sialylation and Nε-lysine protein acetylation. Cells lacking SLC33A1 constitutively trafficked the ATF6α precursor to the Golgi, but exhibit impaired subsequent Golgi processing, correlating with altered ATF6α Golgi glycosylation. SLC33A1 deficiency also deregulated activation of the IRE1 branch of the UPR, pointing to a selective loss of ATF6α-mediated negative feedback in the UPR. Notably, *Slc33a1*-deleted cells accumulated higher levels of unmodified sialylated N-glycans, precursors to acetylated glycans, likely reflecting impaired glycan processing. By contrast, deletion of ER-localised acetyltransferases NAT8 and NAT8B, which catalyse protein Nε-lysine acetylation in the secretory pathway, did not replicate the ATF6α processing defects observed in *Slc33a1*-deficient cells. Together, our findings highlight a role for SLC33A1-mediated metabolite transport in the post-ER maturation of ATF6α and point direct links between small-molecule metabolism and branch-specific signalling in the UPR.

## Introduction

The endoplasmic reticulum (ER) is a central hub for protein folding and the maintenance of cellular homeostasis (Sun & Brodsky, 2019). When misfolded proteins accumulate in the ER, a cellular stress response known as the Unfolded Protein Response (UPR) is triggered to restore protein folding capacity. The UPR is regulated by three known transmembrane sensors: Inositol-Requiring Enzyme 1 (IRE1), Activating Transcription Factor 6 (ATF6), and Protein Kinase R-like ER Kinase (PERK) (Walter & Ron, 2011). In vertebrates, ATF6 plays a pivotal role in restoring ER homeostasis by promoting the expression of molecular chaperones and facilitating ER-associated degradation (ERAD) to eliminate misfolded proteins (Adachi *et al*, 2008). Vertebrates express two isoforms, ATF6α and ATF6β. ATF6α has predominant role in regulating UPR target genes in mammalian cells and is the primary focus of this study. Dysregulation of ATF6α signalling has been implicated in the pathophysiology of neurodegenerative disorders such as Alzheimer’s and Parkinson’s diseases, metabolic diseases, and cancers (Hetz & Papa, 2018). Given its critical function in cellular stress responses, extensive research has been directed toward identifying modulators of ATF6α signalling.

The inactive precursor of ATF6α is an ER resident type II transmembrane protein. In response to ER stress, ATF6α translocates from the ER to the Golgi apparatus (Shen *et al*, 2002). There, it undergoes sequential cleavage by the luminal Site-1 Protease (S1P) and the intra-membrane Site-2 Protease (S2P), releasing a cytosolic fragment (ATF6α-N), which further translocates to the nucleus serving as a transcription factor that activates UPR target genes (Haze *et al*, 1999). N-linked glycosylation of ATF6α in the ER has been implicated in its trafficking and processing (Hong *et al*, 2004), suggesting a potential link between ATF6α activation and the broader network of cellular post-translational modifications involved in regulating its function.

In a recent genome-wide CHO cell-based CRISPR-Cas9 screen for modulators of ATF6α signalling, we identified calreticulin, an ER-resident lectin chaperone, as a selective repressor of ATF6α (Tung *et al*, 2024). By interfering with the conversion of the inactive precursor to the cleaved active form calreticulin maintains ATF6α in an inactive state under physiological conditions. Loss of calreticulin function leads to increased ATF6α activity, suggesting that calreticulin regulates ATF6α and limits UPR activity under homeostatic conditions. This finding emphasises the importance of endogenous repressors of ATF6α in tuning the cellular response to ER stress.

The same CRISPR-Cas9 screen also identified genes whose deletion interfered with ATF6 activation. While some hits, such as genes encoding the proteases known to process the inactive ATF6α precursor into its active form, were anticipated, the discovery that deletion of *Slc33a1* (Solute Carrier Family 33 Member A1), a gene encoding an ER-localised solute transporter, was unexpected highlighting the need for further investigation.

SLC33A1 was first identified in 1997 by Kanamori et al (Kanamori *et al*, 1997). In that study, introducing SLC33A1 into COS-1 cells enabled the formation of O-acetylated gangliosides, suggesting its involvement in the acetylation of sialic acid in the carbohydrate structure of gangliosides. This effect was attributed to the limited capacity of parental COS-1 cells to import acetyl donors into their secretory pathway, alongside the ability of SLC33A1 to import acetyl-CoA (a bulky, charged molecule that cannot freely cross membranes) into the ER lumen. Notably, SLC33A1 displayed characteristics consistent with membrane transporters, such as multiple membrane-spanning domains suggesting a potential role as an ER membrane acetyl-CoA transporter. This proposed function is further supported by a recently published cryo-electron microscopy structure of the human SLC33A1 in complex with acetyl-CoA (Zhou *et al*, 2025).

To date, SLC33A1, also known as AT-1, remains the sole ER-localised transporter identified within cells that facilitates the transport of acetyl-CoA into the ER (Kanamori *et al*, 1997), a process recognised as essential for ER protein N^ε^-lysine acetylation, a well-established reversible post-translational modification of nuclear, cytoplasmic and mitochondrial proteins (Kim *et al*, 2006a). In the secretory pathway, N^ε^-lysine acetylation may influence both the processing of N-linked glycans in the Golgi and the stability of ER-localised proteins (Costantini *et al*, 2007; Choudhary *et al*, 2009; Dieterich *et al*, 2021). Additionally, SLC33A1 has been implicated in providing acetyl-CoA for the acetylation of terminal sialic acid residues during the biogenesis of gangliosides (Zhou *et al*, 2025; Kanamori *et al*, 1997). Therefore, similar to other post-translational modifications, proper acetylation of secretory proteins is a critical step in assisting protein folding and maturation (Jonas *et al*, 2010).

Given its localisation and diverse functional roles, SLC33A1 is well positioned to impact ER and Golgi environments, compartments that are directly involved in ATF6α processing and activation. These roles, along with the broader connections between acetyl-CoA metabolism, ER function, and physiological processes such as aging and neurodegeneration [reviewed in (Brown & Naidoo, 2012; Chandrahas *et al*, 2018; Guo *et al*, 2022)], provided a rationale to explore how SLC33A1 activity might influence ATF6α signalling pathways.

## Results

### CRISPR-Cas9 screening identifies SLC33A1 as novel regulator of ATF6α

To identify genes whose inactivation compromises ATF6α’s stress responsiveness, we used an established CHO-K1 dual UPR reporter cell line harbouring an XBP1s::mCherry and BiP::GFP reporters that monitor IRE1 and ATF6α activity, respectively (Fig. 1A). In a genome-wide CRISPR-Cas9 screen, whose details have been previously published (Tung *et al*, 2024), single-guide RNAs (sgRNA) targeting *Slc33a1* were markedly enriched in ER stressed cells with preserved XBP1::mCherry (IRE1 pathway) activity but diminished BiP::GFP (ATF6α pathway) activity (Fig. 1A and 1B). Notably, five out of six sgRNAs targeting *Slc33a1* in the genome-wide library were progressively enriched through the two rounds of phenotypic selection for stressed cells displaying bright XBP1::mCherry and dull BiP::GFP expression (Fig. 1B). The only sgRNA that was not enriched targeted the 3’end of the gene, likely failing to inactivate it (Supplemental Fig. S1A). These features nominated *Slc33a1* as a candidate gene whose inactivation selectively compromises ATF6α signalling.

**Figure 1.**
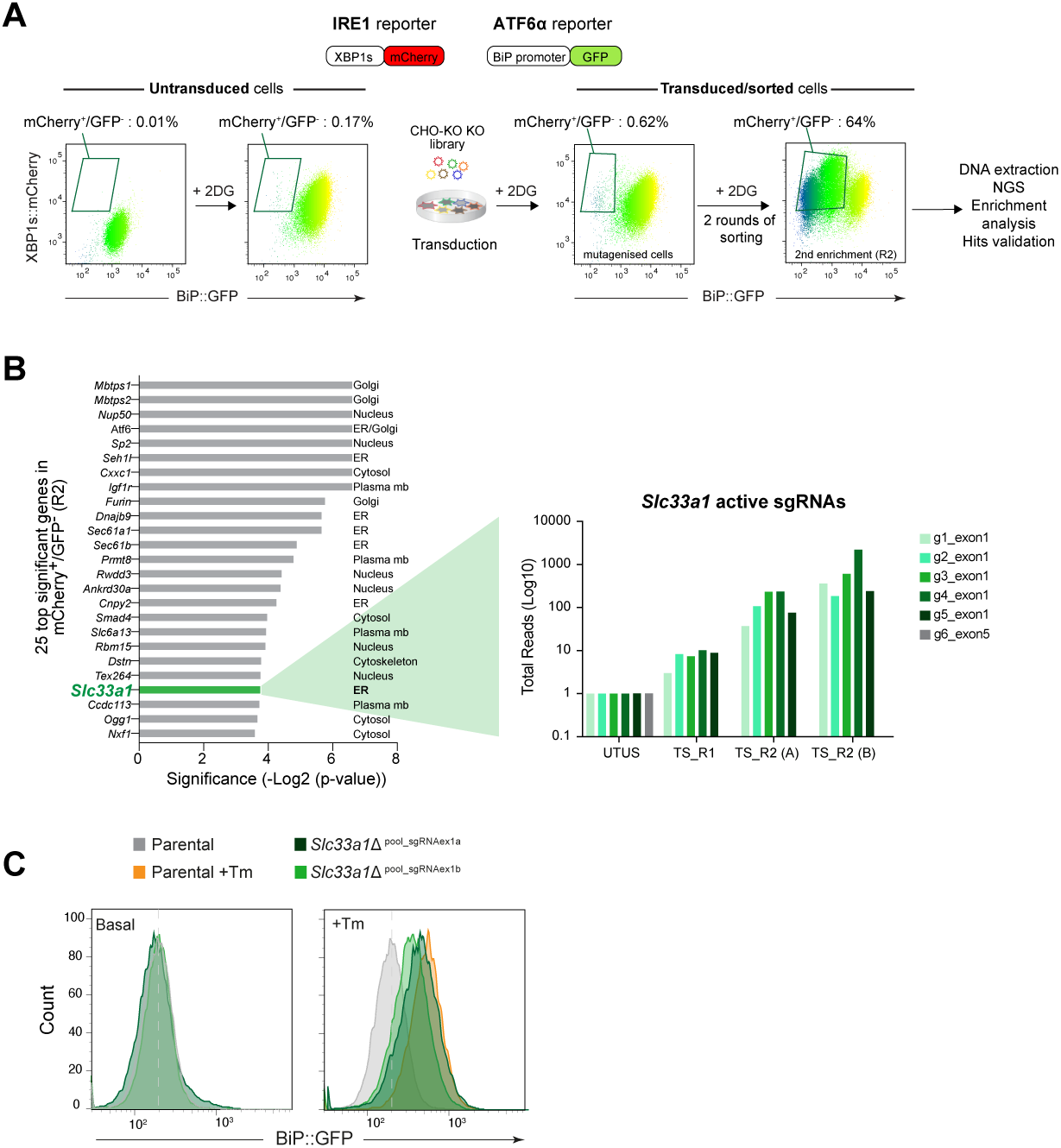
***CRISPR-Cas9 screening identifies SLC33A1 as a positive regulator of ATF6α.* (A)** Schematic overview of a screen for genes whose inactivation by CRISPR-Cas9 selectively compromises the ATF6α branch of the UPR. The screen was performed in dual reporter CHO-K1 cell lines stably expressing XBP1s::mCherry (to monitor IRE1 activity) and BiP::GFP (to monitor ATF6α activity) reporters. Twodimensional contour plots depict the reporter levels. Following transduction with a genome-wide CRISPR knockout (KO) sgRNA library and treatment with 2-deoxy-Dglucose (2DG) to induce ER stress, XBP1s::mCherry-bright; BiP::GFP-dull cells recovered by FACS from the left upper quadrant of the contour plot, were expanded through two rounds of enrichment (R2, sorting). Genomic DNA was extracted for next generation sequencing (NGS), gene ontology enrichment analysis, and validation of candidate genes. **(B)** Left: Significance [-log2 (p-value)] provided for the top 25 genes identified by the criteria noted in ‘1A’. *Slc33a1* gene is highlighted in green. Right: The total read count enrichment of the six sgRNAs targeting *Slc33a1* through the selection process. [UTUS: untreated and unsorted; TS: Treated and sorted; R1: Round 1 of enrichment; R2: Round 2 of enrichment; A-B: pools of cells; g1-g6: sgRNAs]. (C) Histogram of BiP::GFP reporter intensity in parental cells and two derivative *Slc33a1-*deleted (*Slc33a1*Δ) polyclonal pools under basal conditions or following ER stress induced with 2.5 µg/ml Tunicamycin (Tm) for 6 h. A representative dataset from one single experiment is shown.

To confirm the genotype-phenotype relationship suggested by the screen, we targeted the *Slc33a1* locus in CHO-K1 IRE1/ATF6α dual UPR reporter cell line using two distinct CRISPR-Cas9 sgRNAs (Supplemental Fig. S1A). Cell pools were selected based on the expression of a FACS-compatible marker co-expressed with Cas9 and the sgRNA. *Slc33a1*-targeted pooled cells exhibited reduced responsiveness of BiP::GFP to ER stress upon tunicamycin (Tm) treatment (Fig. 1C), consistent with the findings of the screen. These results support the role of *Slc33a1* as a positive modulator of ATF6α-mediated ER stress signalling.

### SLC33A1 depletion has opposing effects on IRE1 and ATF6α signalling

To further investigate the role of SLC33A1 in ATF6α activation, we isolated single clones targeting *Slc33a1* from the pools above and evaluated the genetic lesions by genomic sequencing (Supplemental Fig. S1B). A derivative clone, *Slc33a1*!1^clnA^ (Allele1: *L42Wfs*72*; Allele 2: *L42Gfs*59*) was selected for subsequent studies based on the expected severe disruption of the resulting multipass transmembrane protein, due to the loss of nearly its entire predicted transmembrane helices region (Supplemental Fig. S1C).

As observed in the pool, under basal conditions, the clonal depletion of SLC33A1 did not significantly impact the ATF6α reporter levels (BiP::GFP) when compared to the parental IRE1/ATF6α dual UPR reporter cells (wildtype) (Fig. 2A). However, depletion of SLC33A1 resulted in significant basal de-repression of the IRE1 pathway reporter (Fig. 2A). This was not limited to the reporter: under basal conditions, the clonal *Slc33a1*!1^clnA^ cells had more spliced endogenous XBP1 mRNA (XBP1s) than parental cells (Fig. 2B). Furthermore, disruption of *Slc33a1* in the clonal *Slc33a1*!1^clnA^ cells selectively compromised responsiveness of their BiP::GFP reporter to ER stress while maintaining responsiveness of their XBP1s::mCherry reporter (Fig. 2A and Supplemental Fig. S2A), mirroring the effect observed in pool of *Slc33a1* targeted cells. Similar findings were observed in a second *Slc33a1*-deleted clone (Supplemental Fig. S2B; *Slc33a1*Δ^clnH^).

**Figure 2.**
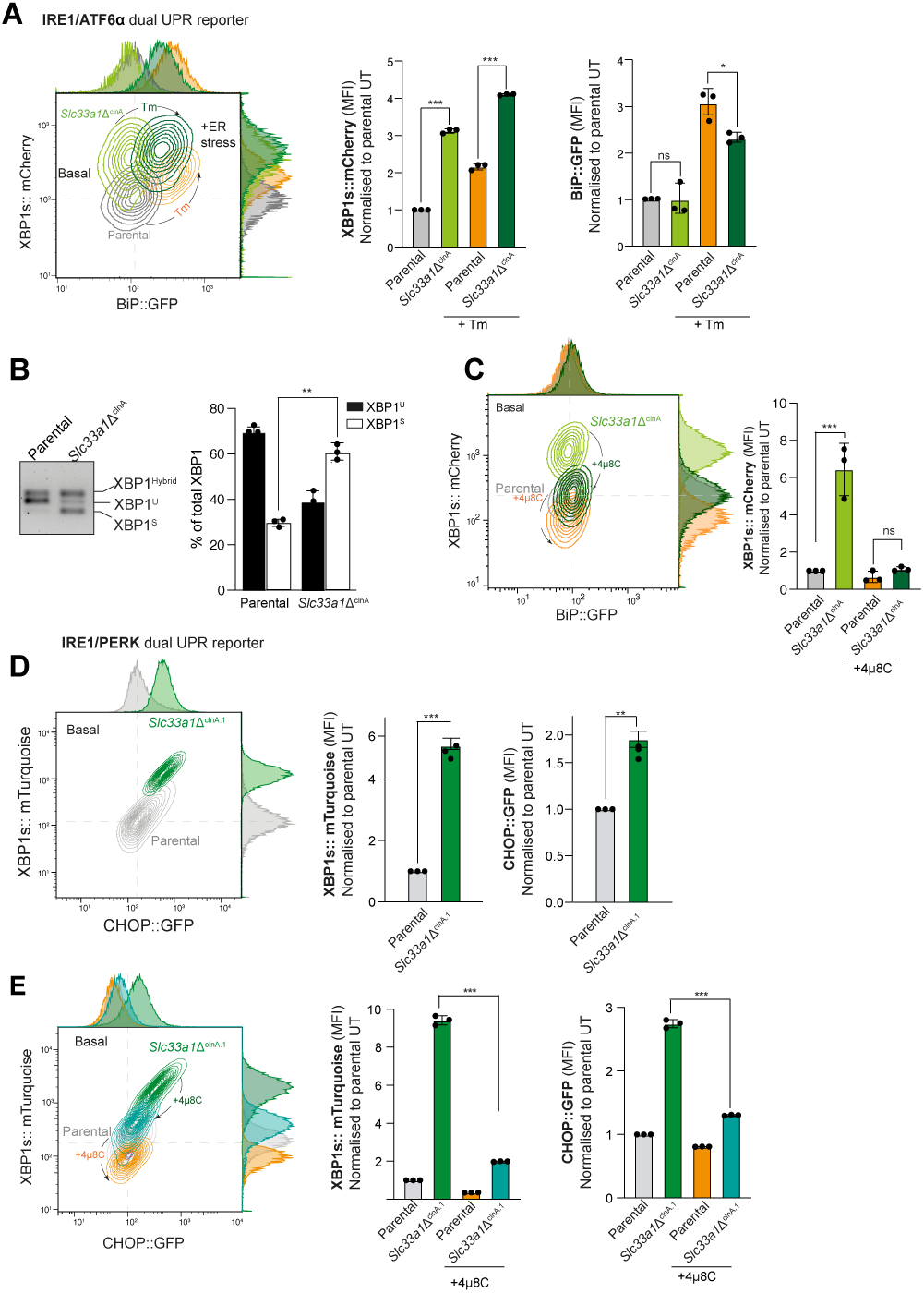
***Depletion of SLC33A1 activates IRE1 and attenuates ATF6α signalling.* (A)** <u>Left:</u> Two-dimensional contour plots of the XBP1s::mCherry and BiP::GFP signals in parental dual UPR reporter cells and a derivative *Slc33a1*-deleted clone (*Slc33a1*Δ^ClnA^), under basal conditions (UT: untreated) (parental in grey; *Slc33a1*Δ^ClnA^ in light green) and after ER stress induction with Tunicamycin (Tm, 2.5 µg/ml, 6 h) (parental in orange; *Slc33a1*Δ^ClnA^ in dark green). <u>Right:</u> Quantification of the median fluorescence intensity (MFI) of XBP1s::mCherry and BiP::GFP in parental and *Slc33a1*Δ^clnA^. Bars represent mean ± SD, and the values of the three replicates as black dots (*p < 0.05; ***p < 0.001, two-sided unpaired Welch’s t-test). **(B)** <u>Left:</u> Representative RT-PCR agarose gel showing XBP1 isoforms in parental and *Slc33a1*Δ^clnA^ cells under basal conditions. The migration of the unspliced (XBP1^U^), spliced XBP1 (XBP1^S^) and hybrid (XBP1^Hybrid^, containing one strand of XBP1^U^ and one strand of XBP1^S^ (Shang & Lehrman, 2004); stained DNA fragments is indicated. <u>Right:</u> Plot of the percentage of XBP1^U^ and XBP1^S^ fractions of XBP1 mRNA in the two genotypes. Bars represent mean ± SD, and the values of the three replicates as black dots (**p < 0.01; two-sided unpaired Welch’s t-test). **C)** <u>Left:</u> Two-dimensional contour plots, as in ‘2A’ of parental and *Slc33a1*Δ^clnA^ dual reporter cells untreated and after 48 h exposure to the IRE1 RNase inhibitor 4µ8C. Untreated: parental in grey; *Slc33a1*Δ^clnA^ in light green, 4µ8C treated: parental in orange; *Slc33a1*Δ^clnA^ in dark green. <u>Right:</u> Quantification of the median fluorescence intensity (MFI) of XBP1s::mCherry in parental and *Slc33a1*Δ^clnA^. Bars represent mean ± SD, and the values of the three replicates as black dots (*p < 0.05; ns: not significant, two-sided unpaired Welch’s t-test). **(D)** <u>Left:</u> Two-dimensional contour plots depicting XBP1s::mTurquoise and CHOP::GFP from parental IRE1/PERK dual UPR reporter parental cells (grey) and a derivate *Slc33a1*-deleted clone (*Slc33a1*Δ^clnA.1^ in green) under basal conditions. <u>Right:</u> Quantification of the median fluorescence intensity (MFI) of XBP1s::mCherry and CHOP::GFP in parental and *Slc33a1*Δ^clnA.1^. Bars represent mean ± SD, and the values of the three replicates as black dots (**p < 0.01; ***p < 0.001, two-sided unpaired Welch’s t-test). **(E)** <u>Left panel:</u> Two-dimensional contour plots, as in ‘2C’ of parental and *Slc33a1*Δ^clnA.1^ dual reporter cells untreated and after 48 h exposure to 4µ8C. Untreated: parental in grey; *Slc33a1*Δ^clnA.1^ in dark green, 4µ8C treated: parental in orange; *Slc33a1*Δ^clnA.1^ in cyan. <u>Right:</u> Quantification of the median fluorescence intensity (MFI) of XBP1s::mCherry and CHOP::GFP in parental and *Slc33a1*Δ^clnA.1^. Bars represent mean ± SD, and the values of the three replicates as black dots (***p < 0.001; two-sided unpaired Welch’s t-test).

To confirm that the elevated XBP1s::mCherry reporter activity observed in the mutant cells was driven by IRE1’s RNase activity, parental and *Slc33a1*-deleted cells were exposed to 4µ8C, an IRE1 RNase inhibitor (Cross *et al*, 2012). Treatment with 4µ8C effectively restored XBP1s::mCherry basal levels in *Slc33a1*-deleted cells, confirming that hyperactivation of the IRE1 pathway observed in SLC33A1 targeted cells proceeded along the conventional pathway (Fig. 2C).

To further explore the role of SLC33A1 in ER homeostasis, we investigated whether depletion of SLC33A1 would affect the PERK branch of the UPR. To this end, we utilised a previously established CHO-K1 reporter cell line stably expressing XBP1s::Turquoise and CHOP::GFP (reporting on the PERK pathway) (Sekine *et al*, 2015). In this background, derivative *Slc33a1*-deleted single clones were generated by CRISPR-Cas9 mutagenesis and a clone, *Slc33a1^cln^*^A.1^, was selected for further studies (Fig. 2D and Supplemental Fig. S1B). The loss of SLC33A1 in the IRE1/PERK dual UPR reporter cell line confirmed the significant de-repression of IRE1 signalling, both by flow cytometry (Fig. 2D) and increased spliced form of the endogenous XBP1 mRNA (XBP1^S^) (Supplemental Fig. S2C), consistent with our findings in the IRE1/ATF6α dual UPR reporter cell line. Loss of SLC33A1 also resulted in constitutive activation of PERK signalling under basal conditions (Fig. 2D), a phenomenon observed in another independently derived *Slc33a1*-deleted clone from the IRE1/PERK reporter cell line (Supplemental Fig. S2D, *Slc33a1*!1^clnE^). Given that IRE1 signalling has been shown to sustain PERK expression (Ong *et al*, 2024), we next assessed whether the constitutive activation of IRE1 could be a contributing factor to the conspicuous upregulation of PERK activity. Indeed, treatment with the selective IRE1 inhibitor 4µ8C restored PERK signalling near to the basal levels (Fig. 2E). These results indicate that the elevated PERK activity observed is, at least in part, a downstream consequence of sustained IRE1 signalling.

### Depletion of SLC33A1 impairs ATF6α processing

The data so far point to a consistent role for SLC33A1 in UPR signalling to enhance IRE1 activity and attenuate ATF6α. However, the potential for cross-pathway negative feedback in the UPR (Kaufman, 2002; Harding *et al*, 2000; Tung *et al*, 2024) leaves open the question of epistasis between the *Slc33a1* mutation and the alterations in UPR signalling. To address this, we investigated the impact of SLC33A1 on ATF6α signalling in further detail. Considering the critical role of ATF6α processing by site-1 (S1P) and site-2 (S2P) Golgi proteases for the generation and trafficking of the active N-terminal domain of ATF6α (N-ATF6α) from Golgi apparatus to nucleus (Haze *et al*, 1999), we next investigated the impact of the loss of SLC33A1 on ATF6α processing by immunoblotting. To this end, we turned to a previously characterised ATF6α knock-in clone, in which the protein encoded by endogenous *Atf6α* locus of CHO-K1 cells was tagged with 3×FLAG-mGreen Lantern (mGL) (Fig. 3A, upper panel) (Tung *et al*, 2024). In this background, we targeted the *Slc33a1* locus and identified a mutant clone (*Slc33a1*Δ^clnA9^) that had early frame-shifts in both alleles and a phenotype of basally elevated endogenous XBP1 splicing (Fig. 3A, lower panel and Supplemental Fig. S1B), consistent with that observed in previously characterised *Slc33a1* mutant clones (see Fig. 2B and Supplemental Fig. S2C).

**Figure 3.**
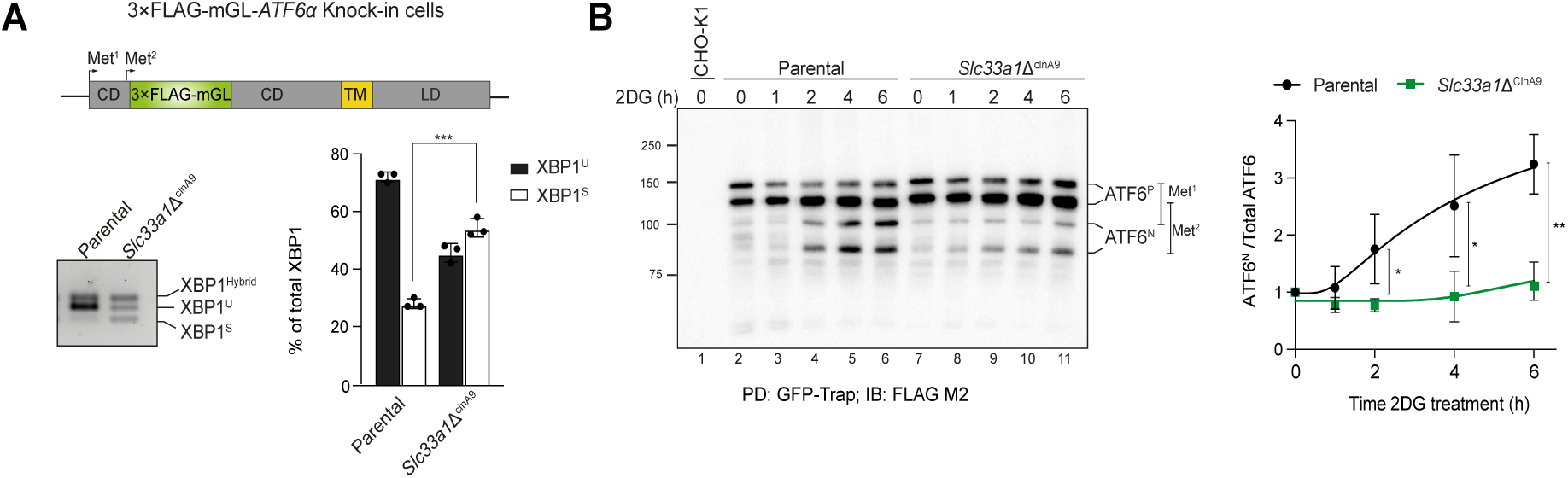
***SLC33A1 deletion impairs ATF6α processing. (*A)** <u>Top:</u> Schema of the genetically modified endogenous CHO-K1 *Atf*6α locus, encoding ATF6α tagged with a 3×FLAG-mGreen Lantern (mGL) upstream of the second methionine (Met) start codon. The first methionine in exon 1 (Met^1^) is also marked. Colour coding: 3×FLAG-mGL (green), cytosolic domain (CD) (grey), transmembrane domain (TM) (yellow), and luminal domain (LD) (grey). <u>Bottom:</u> Representative agarose gel showing XBP1 cDNA isoforms detected via RT-PCR in 3×FLAG-mGL-ATF6α knock-in parental cells and *Slc33a1*Δ^clnA9^ cells under basal conditions. The migration of the unspliced (XBP1^U^), spliced XBP1 (XBP1^S^) and hybrid (XBP1^Hybrid^, containing one strand of XBP1^U^ and one strand of XBP1^S^ (Shang & Lehrman, 2004); stained DNA fragments is indicated. Plot of the percentage of XBP1^U^ and XBP1^S^ fractions of XBP1 mRNA in the two genotypes. Bars represent mean ± SD, and the values of the three replicates as black dots (**p < 0.01; two-sided unpaired Welch’s t-test). **(B)** <u>Left:</u> Anti-FLAG M2 immunoblot of 3×FLAG-mGL-tagged ATF6α in lysates of parental and mutant *Slc33a1*Δ^clnA9^ cells exposed to 2-deoxy-D-glucose, (2DG, to induce ER stress) for the indicated period of time. The two forms of full-length ATF6α precursor (ATF6^P^) and two processed N-terminal ATF6α forms (ATF6^N^), arising from alternative translational initiation sites (Met^1^ or Met^2^), are indicated. Parental CHO-K1 untagged cells (line 1) were used as control for background. The immunoblot is representative of three experiments. <u>Right:</u> Ratio of the signal from both processed ATF6^N^ forms to the total ATF6α signal (ATF6^N^ + ATF6^P^) for each genotype. Data are presented as the mean ± SD from three independent experiments. *p < 0.05 and **p < 0.01, according to 2way ANOVA test followed by a Bonferroni *post hoc* test. PD indicates pull-down, while IB refers to immunoblot.

The knock-in *Slc33a1*Δ^clnA9^ cells were treated with the mild ER stress causing agent 2-deoxy-D-glucose (2DG) for various time intervals, and 3×FLAG-mGL-ATF6α was purified using GFP-trap Agarose (ChromoTek), followed by anti-FLAG immunoblotting. Four distinct bands were observed, likely due to the presence of two methionine residues that can serve as alternative translation initiation sites in the tagged mRNA encoded by the knock-in ATF6α allele (Fig. 3A, upper panel). This results in the production of two precursors ATF6α isoforms (ATF6^P^) with different lengths and their corresponding two processed N-terminal ATF6α fragments (ATF6^N^). The ratio of processed N-ATF6α forms (ATF6^N^) to total ATF6α was quantified to assess ATF6α processing under ER stress (Fig. 3B). *Slc33a1*-deleted cells exhibited consistently reduced levels of the processed N-ATF6α form upon ER stress, compared to the parental cells. These findings align with our previous flow cytometry observations and suggest that SCL33A1 positively contributes to ATF6α processing.

### Constitutive Golgi localisation of ATF6α in *Slc33a1Δ* cells

Stress-dependent translocation of the membrane-bound ATF6α precursor from the ER to the Golgi is a pre-requisite for its proteolytic activation (Chen *et al*, 2002). To determine whether defective ATF6α processing in cells lacking SLC33A1 correlates with defective trafficking, we used 3×FLAG-mGL-ATF6α knock-in parental cells and their *Slc33a1*-inactivated counterpart (*Slc33a1*Δ^clnA9^) for live-cell microscopy (Fig. 4A). Under ER stress, parental cells exhibited a conspicuous increase in nuclear mGL-ATF6α signal compared to basal conditions that was absent in stressed *Slc33a1*Δ^clnA9^ cells (Fig. 4B and 4C). This observation aligned with the reduced processed N-ATF6α form observed in *Slc33a1*Δ cells on immunoblot (Fig. 3B).

**Figure 4.**
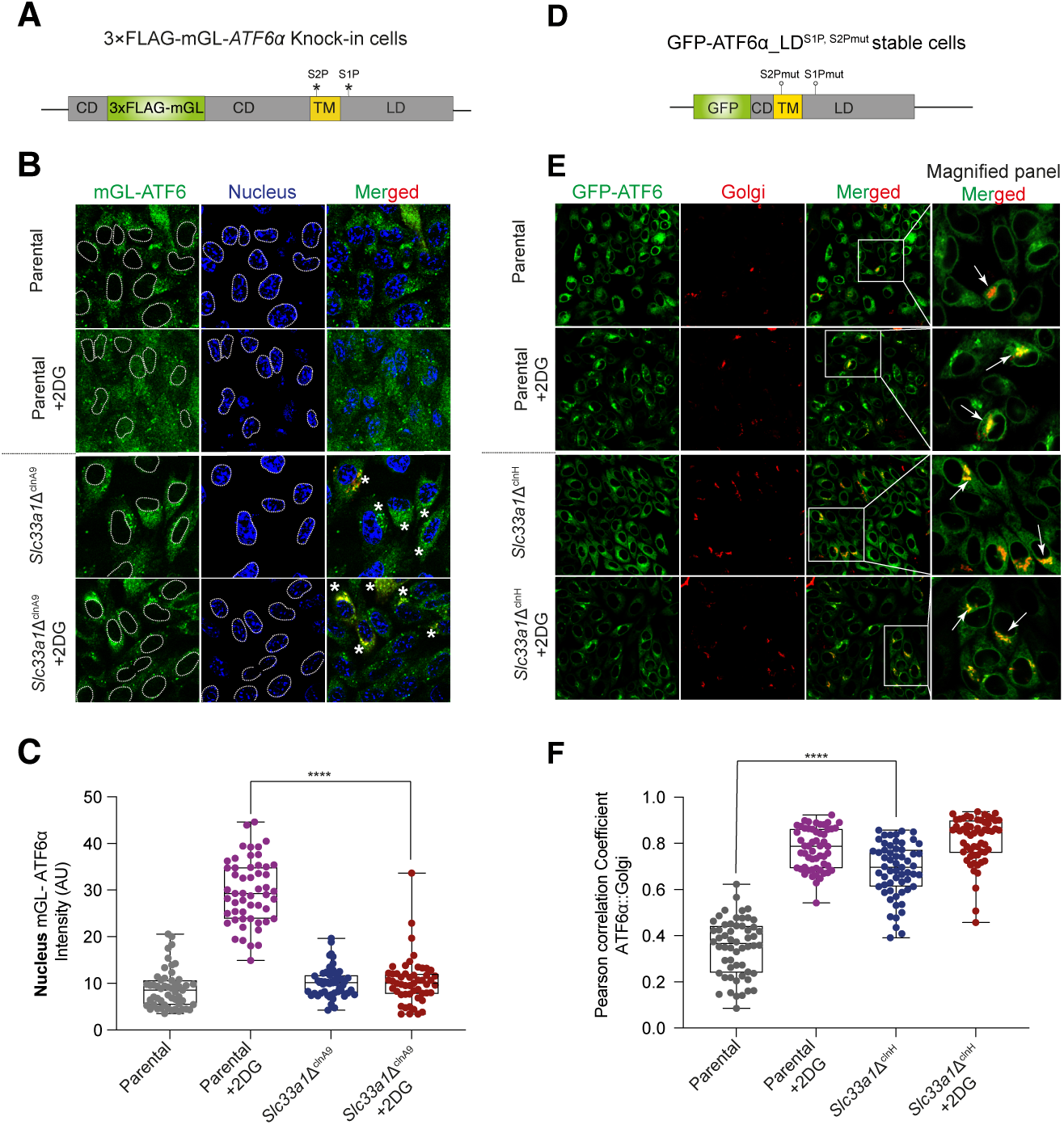
***Constitutive Golgi localisation of ATF6α in SLC33A1Δ CHO cells.* (A)** Schema of the genetically modified endogenous *Atf6α* locus (as in Fig. 3A). The S1P and S2P cleavage sites are indicated by stars. **(B)** Representative live-cell confocal microscopy images of 3×FLAG-mGL-ATF6α knock-in parental cells and *Slc33a1*Δ^clnA9^ derivative clone under basal and ER stress conditions (+2DG). mGL-ATF6α is shown in green, and the nucleus is in blue. Cells were transiently transfected with the pmScarlet_Giantin-C1 plasmid to label the Golgi apparatus (see merged panel). Co-localisation of mGL-ATF6α to a punctate pattern (Golgi-like, yellow) is indicated by asterisks in the merged panel. Scale bar: 20 μm. **(C)** Quantification of mGL-ATF6α signal intensity in the nucleus (outlined by dashed lines), measured using ImageJ software from images in panel ‘4B’. Data from parental cells (n > 50) and *Slc33a1*Δ^clnA9^ cells (n > 50) are shown as a box-and-whisker plot, displaying all individual values, including minimum and maximum intensities. Statistical significance was determined using a two-sided unpaired Welch’s t-test (p**** < 0.0001). **(D)** Schema of the stable transgene encoding an N-terminally tagged ATF6α derivative (as in ‘4A’ above) in which GFP replaces most of the cytosolic domain, and mutations in the S1P and S2P cleavage sites preclude activating proteolysis and favour Golgi retention of trafficked protein. Colour coding: GFP tag (green), cytosolic domain (CD, grey), transmembrane domain (TM, yellow), and luminal domain (LD, grey). **(E)** Representative live-cell confocal microscopy images as in ‘4B’ above. Cells were transiently transfected with pmScarlet_Giantin-C1 to visualise the Golgi (in red). Insets show magnified regions, with arrows highlighting GFP-ATF6α co-localisation with the Giantin Golgi marker. Scale bar: 20 μm. **(F)** Pearson’s correlation coefficients were used to quantify co-localisation of GFP-ATF6α_LD^S1P,S2Pmut^ with Giantin in parental cells (n > 50) and *Slc33a1*Δ^clnH^ cells (n > 50). Data are presented as a box-and-whisker plot, including all individual values along with minimum and maximum intensities. Co-localisation analysis was performed using Volocity software, and statistical significance was determined using a two-sided unpaired Welch’s t-test (****p < 0.0001).

Interestingly, the defect in ATF6α processing and nuclear accumulation of the processed N-terminal 3×FLAG-mGL-ATF6α in *Slc33a1*Δ^clnA9^ cells was not associated with retained signal in the ER membrane, but rather with an enhanced punctate signal that co-localised (or overlapped) with a Golgi marker (Fig. 4B, asterisks). This feature suggests that SLC33A1 acts at a downstream step after ATF6α trafficking to the Golgi.

Resistance to proteolytic cleavage by S1P and S2P uncouples ER to Golgi trafficking from the downstream processing of ATF6α and facilitating interrogation of the two events separately. Therefore, we used a previously described CHO-K1 cell stably expressing a GFP-tagged version of cgATF6α_LD (luminal domain), with mutations in the S1P and S2P cleavage sites (GFP-ATF6α_LD^S1P,S2Pmut^) (Tung *et al*, 2024) to follow up on the suggestion of a perturbation to ATF6α’s itinerary in cells lacking SLC33A1 (Fig. 4D) (Tung *et al*, 2024). A derivative *Slc33a1*-deleted clonal cell (called *Slc33a1*Δ^clnH^) was generated for analysis (Supplemental Fig. S1B). Both under basal and stressed conditions the GFP-ATF6α_LD^S1P,S2Pmut^ signal in the *Slc33a1* mutant cells was distinctly punctate with significant co-localisation to a Golgi marker (Fig. 4E and 4F). In contrast, parental cells exhibited a diffuse GFP-ATF6α signal under basal conditions, with noticeable co-localisation of GFP-ATF6α with the Golgi marker only upon exposure to ER stress (Fig. 4E and 4F). This observation was reproduced in a second *Slc33a1*-deficient GFP-ATF6α_LD^S1P,S2Pmut^ clone (Supplement Fig. S3, *Slc33a1*Δ^clnK^).

The punctate Golgi-associated localisation of the fluorescent ATF6α-fusion proteins under basal conditions is consistent with disruption of an ATF6α-driven feed-back loop in the *Slc33a1*Δ^clnA9^ and *Slc33a1*Δ^clnH^ cells that functions homeostatically, even in the absence of pharmacologically imposed stress. These findings suggest that in the absence of SLC33A1, ATF6α constitutively translocates to the Golgi and encounter a block in its down-stream processing.

### Impaired Golgi-type sugar modification of ATF6α in *Slc33a1Δ* cells

Upon arrival, the glycans on the luminal domain of the ATF6α precursor become substrates for Golgi localised enzymes. Given the evidence above for perturbation of a Golgi-localised node in ATF6α’s itinerary and the implication of SLC33A1 in providing the acetyl-CoA building blocks for post-ER glycan modification, we wished to compare the maturation of ATF6α glycans in parental and *Slc33a1*-mutant cells.

ATF6 contains three evolutionary conserved N-linked glycosylation sites in its ER luminal domain, and its N-glycosylation state has been suggested to influence its trafficking (Hong *et al*, 2004). O-linked glycosylation of ATF6 has also been reported (Shen *et al*, 2002), although the functional implications of these Golgi-modified sugars remain unclear. To reveal the palette of ATF6 glycosylation states, we introduced the GFP-tagged version of cgATF6α_LD (lacking the S1P and S2P cleavage sites; GFP-ATF6α_LD^S1P, S2Pmut^) into CHO-K1 and analysed the mobility of GFP-tagged proteins by autoradiography on SDS-PAGE of ^35^S-metabolically labelled material immunopurified from stressed cells. Cells lysates were subjected to enzymatic digestion with different glycosidases to characterise ER-and Golgi-glycosylation modifications (Fig. 5A and 5B). In the absence of enzymatic digestion, four distinct ATF6α-specific bands appeared (Fig. 5B, Lane 1). The three lower molecular weight bands were labelled as 1N-3N, denoting the number of N-glycans attached to ATF6α. A fourth, higher molecular weight band was interpreted as a mixture of complex-type glycans (CG) and O-glycans (OG) forms of ATF6α (labelled CG+OG), typically generated in the Golgi. As expected, upon treatment with Endo H (Fig. 5B, Lane 2), the N3, N2, and N1 bands collapsed into a single, faster-migrating band, indicating that these represent simple N-glycans characteristic of the ER-localised form of ATF6α. The upper CG+OG band remained Endo H-resistant, consistent with the presence of Golgi-modified glycans. When treated with PNGase F alone, which cleaves both N-glycans and CG forms (Fig. 5B, Lane 3), N3 and N2 collapsed to N1 (as expected). The upper CG+OG band shifted to a slightly slower-migrating species (labelled as OG), suggesting a mixed composition of N-and O-glycans. This was confirmed by its disappearance from samples co-treated with O-glycosidases (Fig. 5B, Lane 4 and 5).

**Figure 5.**
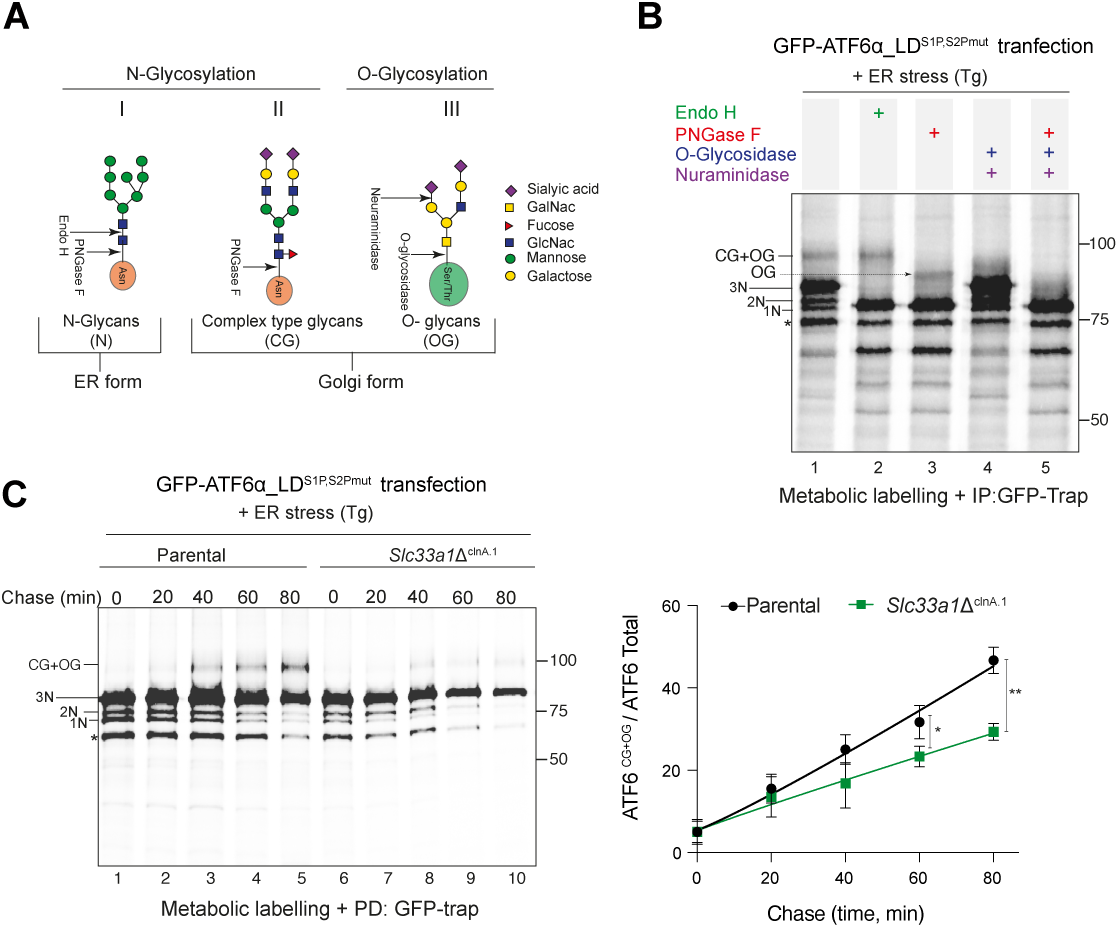
***Impaired Golgi-type sugar modification of ATF6α in Slc33a1Δ cells.* (A)** Schema of glycan modifications in the ER and Golgi and their sensitivity to glycosidases. **(B)** Autoradiograph of GFPATF6α_LDS1P,S2Pmut immunoprecipitated from cell lysates of 35S-Met/Cysmetabolically-labelled transfected CHO-K1 cells. To induce ATF6α trafficking from ER to Golgi, cells were treated with Thapsigargin (Tg, 300 nM). Samples were exposed to the indicated glycosidases. The bands corresponding to ATF6α-LD (luminal domain) with one (1N), two (2N), or three (3N) N-glycan modifications are indicated. O-glycan and complex-type glycans are annotated as OG and CG, respectively. (*) denotes a non-specific band. **(C)** Left: Autoradiograph as in ‘5B’ of samples from a pulse-chase experiment in cells of the indicated genotype. Cells were treated with Tg for 30 min before the pulse, and Tg treatment was maintained during the 3⁵S Met/Cys pulse labelling (15 min) and the chases times (20-80 min). Shown is a representative of three experiments. <u>Right:</u> Ratio of the signal from complex-type (CG) and O-glycan (OG)-modified GFP-ATF6α-LD^S1P,S2Pmut^ to the total ATF6α signal for each genotype over the chase period. Data are presented as the mean ± SD, n = 3; *p < 0.05 and **p < 0.01, according to 2way ANOVA test followed by a Bonferroni *post hoc* test.

With figure 5B as a reference, we compared the time-dependent changes in glycosylation of the GFP-ATF6α_LD^S1P,S2Pmut^ sentinel in parental cells and the *Slc33a1*Δ^ClnA.1^ derivative clone by pulse-chase labelling under ER stress conditions. In parental cells, there was a progressive increase in the CG+OG Golgi-modified ATF6α population over the chase period (Fig. 5C, lanes 3-5). Interestingly, in *Slc33a1*Δ cells, the formation of these Golgi-modified populations were significantly reduced (Fig. 5C, lanes 8-10).

Taken together our findings point to an uncoupling between trafficking to the Golgi, which is enhanced in *Slc33a1* mutant cells, and Golgi modification of glycans and proteolytic processing of ATF6α by S1P and S2P Golgi proteases, which is diminished by the depletion of SLC33A1, to attenuate subsequent translocation to the nucleus and ultimately ATF6α signalling.

O-acetylation of sialic acids is a post-translational modification that modulates the stability, trafficking, and function of secreted glycoproteins (Varki, 2008). In the Golgi, acetyl groups from acetyl-CoA are transferred to sialic acid residues at positions C4, C7, C8, or C9 on nascent glycoproteins (Arming *et al*, 2011). This modification protects sialic acids from sialidase-mediated cleavage, preserves glycan integrity during Golgi processing, and can influence glycoprotein sorting by altering recognition by lectins and cargo receptors. Since SLC33A1 has been reported to play a role in ganglioside formation by supplying acetyl-CoA to the Golgi for O-acetylation, we sought to detect O-acetyl modification of ATF6α in wild-type and *Slc33a1-*deleted cells by mass spectrometry. While we successfully identified terminal sialic acid modifications on the N-glycans of ATF6α, we were unable to detect their O-acetylated forms, likely due to the inherent technical limitations of mass spectrometry in capturing these labile modifications, which are prone to loss during ionization and fragmentation. However, we observed a marked increase in the unmodified sialylated N-glycans, the precursors to O-acetylated forms, in the *Slc33a1*-deleted cells compared to the parental cell line. This finding is consistent with accumulation of sialylated glycans that remain unacetylated due to defective glycan processing in the absence of SLC33A1 (Supplemental Fig. S4).

### Depletion of acetyl-CoA acetyltransferases does not phenocopy SLC33A1 loss

SLC33A1 has been reported as an acetyl-CoA transporter, facilitating the import of acetyl-CoA from the cytosol into the ER lumen and the Golgi apparatus via vesicular transport (Kanamori *et al*, 1997; Ko & Puglielli, 2009; Hirabayashi *et al*, 2013). Within the ER lumen, acetyl-CoA has been proposed to serve as a substrate for the ER membrane-bound lysine acetyltransferases of NAT8 family of enzymes (Pehar & Puglielli, 2013) (Fig. 6A). These acetyltransferases transfer the acetyl group from acetyl-CoA to N^ε^-lysine of various protein substrates, a process that is thought to affect protein secretion, molecular stabilisation, conformational protein assembly and overall cellular homeostasis (Kouzarides, 2000; Farrugia & Puglielli, 2018; Peng & Puglielli, 2016). To determine whether defective protein N^ε^-lysine acetylation contributes to the phenotype observed in *Slc33a1-*deleted cells, we measured the consequences of *Nat8* inactivation on ATF6α signalling.

**Figure 6.**
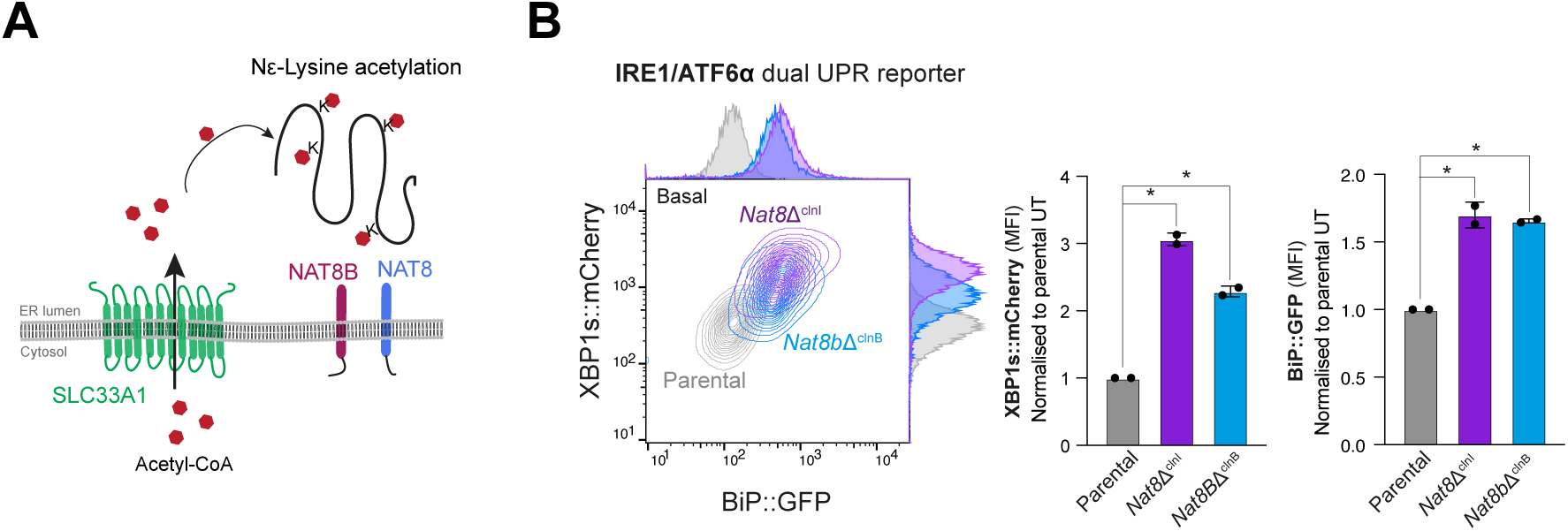
***Depletion of ER-localised acetyltransferases does not phenocopy SLC33A1 loss.* (A)** Schematic representation of N^ε^-lysine acetylation in the ER. Acetyl-CoA (red hexagons), transported into the ER lumen of the secretory pathway by SLC33A1, serves as a acetyl donor for NAT8 and NAT8B, two luminal acetyl-transferases that modify lysine residues on protein substrates, resulting in N^ε^-lysine acetylation. **(B)** <u>Left</u> <u>panel:</u> Two-dimensional contour plots depicting XBP1s::mCherry and BiP::GFP signals in parental IRE1/ATF6α dual UPR reporter cells (grey) and *Nat8*-deleted derivative clones (*Nat8*Δ^clnI^: purple; *Nat8b*Δ^clnB^: blue). <u>Right</u> <u>panel</u>: Quantification of the median fluorescence intensity (MFI) of XBP1s::mCherry and BiP::GFP from two independent experiments is shown in bar diagrams representing mean ± SD. Statistical analysis was performed using a two-sided unpaired Welch’s t-test (*p < 0.05).

CHO-K1 cells express two isoforms of the *Nat8* gene: *Nat8* and *Nat8b*, redundancy amongst which could explain failure to detect them in genome-wide screens. Both isoforms were targeted independently by CRISPR-Cas9 gene editing in IRE1/ATF6α dual UPR reporter cell lines to generate derivative *Nat8*-deleted clones. The consequences of *Nat8* depletion were assessed by measuring reporter activity via flow cytometry. Targeting both *Nat8* isoforms did not replicate the phenotype observed in *Slc33a1*-deleted cells. Instead, inactivation of either *Nat8* or *Nat8b* resulted in elevated reporter activity of both XBP1::mCherry and BiP::GFP (Fig. 6B). These findings support a role for N^ε^-lysine acetylation in maintaining ER protein homeostasis and suggest that *Nat8* and *Slc33a1* influence ER homeostasis and ATF6 signalling through distinct mechanisms.

## Discussion

This unbiased genome wide CRISPR-Cas9 screen establishes a link between the ER localised transmembrane solute transporter SLC33A1 and ATF6α activation. Initially established by tracking the activity of a UPR pathway reporter, the role of *Slc33a1* as a positive and selective regulator of ATF6α activity was confirmed by gene deletion and examining endogenous UPR markers. The defect caused by the deletion of *Slc33a1* mapped to a post-ER step in the itinerary of ATF6α processing. It correlated with a defect in activating-proteolysis of the ATF6α precursor and in post-ER processing of its glycans, despite its constitutive trafficking from the ER to the Golgi. These findings point to an unanticipated role for ER metabolite(s) in promoting ATF6α signalling.

Our work highlights that *Slc33a1*’s deletion has discordant effects on UPR pathways: they promote IRE1 activity while inhibit ATF6α activity. This pattern could reflect pleiotropic effects of SLC33A1 on distinct branches of the UPR [and more broadly on cellular adaptations to stress (Pehar *et al*, 2012)], but it is insufficient to discriminate primary from secondary effects. However, considering the known cross-talk between UPR branches, the impaired processing of Golgi-localised ATF6α offers a parsimonious explanation for the observations reported here: ATF6α’s dominant role in regulating chaperone gene expression in the vertebrate UPR (Yamamoto *et al*, 2007; Ishikawa *et al*, 2011) specifies disruption of ER proteostasis, de-repressing both IRE1 signalling and the upstream-most steps in ATF6α activation in *Slc33a1* mutant cells. This model is consistent with the enhanced splicing of XBP1 and the constitutive trafficking of the ATF6α precursor to the Golgi observed in the *Slc33a1* mutant cells. The smooth transit of ATF6α from the ER to the Golgi is consistent with findings that point to normal ER to Golgi trafficking of a model secretory protein in *Slc33a1* mutant mouse cells (Dieterich *et al*, 2021). However, the block in ATF6α processing once arrived in the Golgi represents an unanticipated finding.

The genetic analysis described here outlines a plausible picture of the epistasis between *Slc33a1*, *Atf6α* and *Ern1* (encoding IRE1), but does not yet uncover the underlying molecular mechanism. While phenotypic data from humans (Huppke *et al*, 2012) and mice (Cooley *et al*, 2021; Dieterich *et al*, 2021) with *Slc33a1* mutations are consistent with secretory pathway dysfunction, they do not currently clarify how SLC33A1 affects ATF6α processing and field is therefore wide open for speculation. It is important to acknowledge that, while the evidence supporting acetyl-CoA transport by SLC33A1 is strong, there is nothing to argue against the possibility that other acetyl-CoA-related ligands containing a 3′-phosphorylated ADP moiety, such as ATP, dATP, and ADP, which have recently been shown to be transported by SLC33A1 (Zhou *et al*, 2025), might also contribute to defective ATF6α activation observed in *Slc33a1* mutant cells. This aligns with ongoing questions about how metabolites like ATP, known to play a key role in protein folding in the secretory pathway (Chaudhry *et al*, 2003; Chakravarthi *et al*, 2006), are imported into the ER.

Nonetheless, the prism of defective acetyl-CoA transport into the ER offers valuable insight. The divergent phenotypes of *Nat8* and *Slc33a1* inactivation argue against a central role for Nε-lysine acetylation in the observed ATF6α activation defect, though this must be interpreted with caution due to potential differences in mutation strength and pathway-specific pleiotropy.

O-acetylation of terminal sialic acid residues on N-Glycans, which relies on a supply of acetyl-CoA as a donor, is believed to be a trans-Golgi modification (Bhide & Colley, 2017), whereas activating processing of the ATF6α precursor is believed to occur in the cis-Golgi (Shen & Prywes, 2004). Nonetheless, it remains possible that a so far unrecognised acetyl-CoA-dependent modification promotes ATF6α processing. Alternatively, given the evidence for a broad defect in secretion in *Slc33a*1 compromised cells, it is equally plausible that a deficiency in acetyl-CoA import (or some other metabolite) results in a widespread impairment of Golgi function, which, among other consequences, disrupts ATF6α activation (Fig. 7).

**Figure 7.**
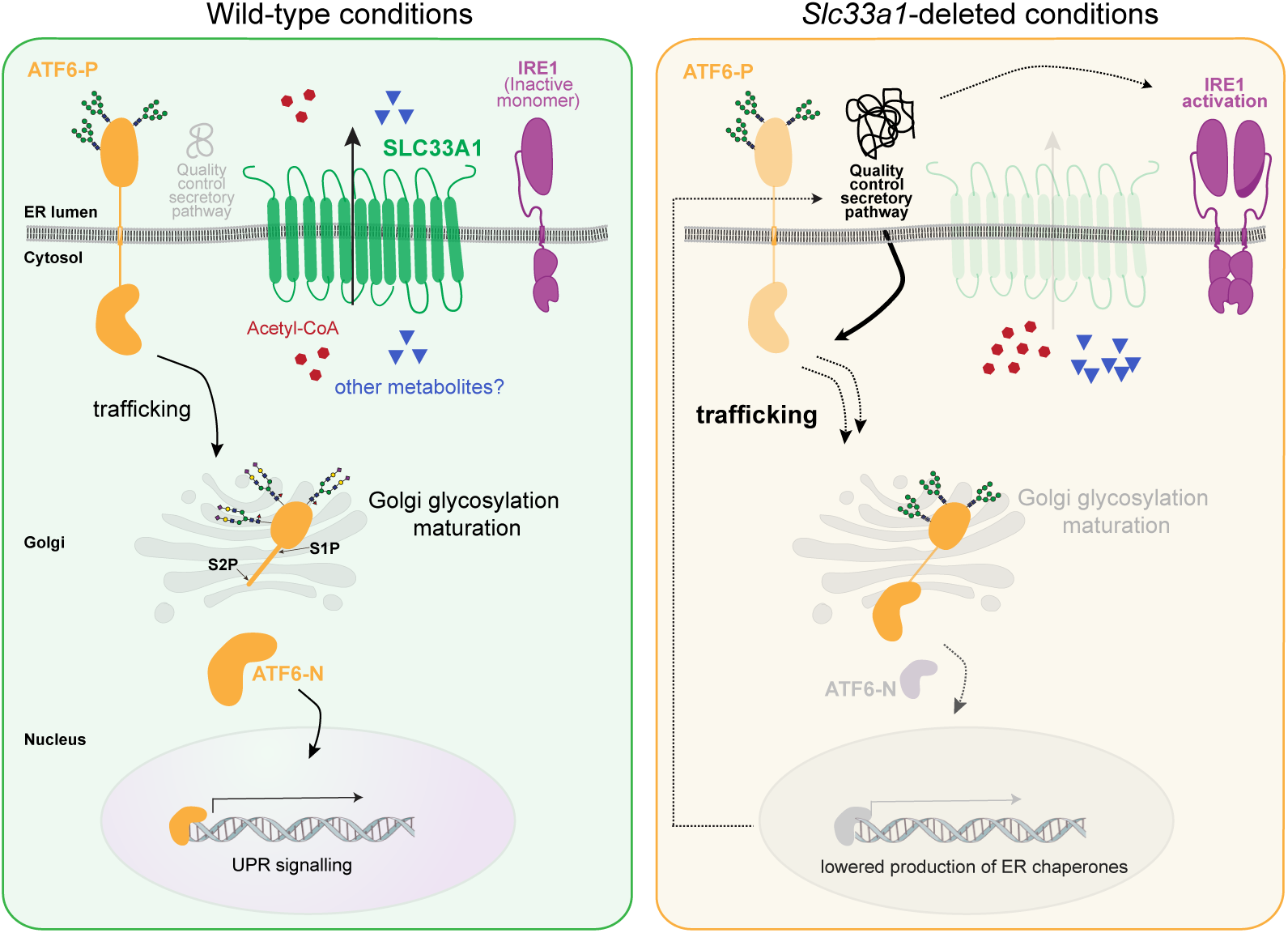
***Proposed model.*** SLC33A1, a multipass transmembrane protein located at the endoplasmic reticulum (ER), transports acetyl-CoA from the cytosol into the ER, to serve as a substrate for acetylation reactions in the secretory pathway. Changes in metabolite abundance caused by loss of SLC33A1 (right panel) impair the processing of the transmembrane ATF6 precursor (ATF6-P) in the Golgi and the liberation of its active fragment, ATF6-N, by the site-specific proteases S1P and S2P, a defect that correlates with altered glycosylation of ATF6-P. Lower ATF6α signalling in the nucleus reduces production of ER chaperones and subsequently impairs protein folding homeostasis in the ER. The resulting enhanced burden of unfolded proteins explains both the heightened IRE1 activity and the constitutive trafficking of ATF6 to the Golgi observed in cells lacking *Slc33a1*. SLC33A1 activity may impinge on other aspects of cellular function. While SLC33A1 may also affect other cellular processes or transport additional metabolites, the chain of events uncovered here attributes a selective role to metabolites transported into the secretory pathway in ATF6 activation.

Whether defective ATF6α activation contributes to the clinical manifestations of SLC33A1 mutations remains to be determined. However, given emerging links between intermediary metabolism, aging and proteostasis in the secretory pathway, this work underscores SLC33A1 as a key genetic entry point for dissecting how metabolic state regulates the unfolded protein response and serve as a stimulus for further research.

## Material and Methods

### Mammalian cell culture

Adherent Chinese Hamster Ovary (CHO-K1) cells (ATCC CCL-61) were cultured in Ham’s F12 nutrient mixture (Sigma), supplemented with 10% (v/v) serum (FetalClone-2, HyClone), 2 mM L-glutamine (Sigma), and 1% penicillin/streptomycin (Sigma). The cells were incubated at 37°C in a humidified environment containing 5% CO2. As specified, cells were exposed to Tunicamycin (Tm, 2.5 µg/ml), 2-deoxy-D-glucose (2DG, 4 mM), 4µ8C (4 µM), dithiothreitol (DTT, 2 mM) or Thapsigargin (Tg, 300nM) for varying times depending on the specific experiments as indicated. Untreated cells were treated with dimethyl sulfoxide (DMSO) solvent vehicle control. Transfections in CHO-K1 cells were performed using Lipofectamine LTX (Thermo Fisher Scientific, USA) at 1:3 DNA (µg) to LTX (µl) ratio.

### Flow cytometry analysis

To assess UPR reporter activity, cells were seeded in 6-well plates and grown to approximately 80–90% confluency before being treated with the specified compounds for the indicated period. For flow cytometry analysis, cells were washed twice with PBS then collected in PBS containing 4 mM EDTA and analysed using a BD LSR Fortessa flow cytometer (BD Biosciences), with 20,000 cells processed per sample. For FACS sorting, cells were harvested in PBS containing 4 mM EDTA and 0.5% bovine serum albumin (BSA) and sorted using a Beckman Coulter MoFlo cell sorter. Sorted cells were either collected in fresh culture medium containing 20% (v/v) serum (FetalClone-2, HyClone), as bulk populations or individually deposited into 96-well plates for subsequent expansion. Live cells were gated based on FSC-A/SSC-A profiles, while singlets were gated using FSC-W/SSC-A. Reporter fluorescence was detected as follows: BiP::GFP using a 488 nm excitation laser and a 530/30 nm emission filter; XBP1s::mCherry using a 561 nm excitation laser and a 610/20 nm emission filter; mTurquoise and BFP using a 405 nm excitation laser and a 450/50 nm emission filter. FlowJo v10 (BD Biosciences) was used for data analysis, and median fluorescence intensity was further analysed and plotted using GraphPad Prism 10.

### RNA extraction and cDNA synthesis

Total RNA was extracted using TRIzol™ Reagent (Invitrogen), with cells incubated in the reagent for 10 min before transferring to clean tubes. To each sample, 200 µl of chloroform was added, followed by vigorous vortexing for 1 min and centrifugation at 13,500 × g for 15 min at 4°C. The upper aqueous phase was carefully removed and mixed with an equal volume of 70% ethanol. RNA was then purified using the PureLink™ RNA Mini Kit (Invitrogen) according to the manufacturer’s protocol and eluted for downstream use. RNA concentrations were measured using a NanoDrop spectrophotometer, and the integrity of 2 µg RNA from each sample was checked on a 1.2% agarose gel.

For reverse transcription PCR (RT-PCR), 2 µg of total RNA was denatured at 70°C for 10 min in a reverse transcription buffer (Thermo Scientific; Cat #EP0441) containing 0.5 mM dNTPs and 0.05 mM Oligo(dT)18 primers. The reaction was then supplemented with 0.5 µl of RevertAid Reverse Transcriptase (Thermo Scientific) and 100 mM DTT and incubated at 42°C for 90 min. The synthesized cDNA was diluted 1:4 for further analysis.

### PCR-based analysis of XBP1 mRNA splicing

Spliced (XBP1s) and unspliced (XBP1u) variants of XBP1 mRNA were amplified from synthesized cDNA using PCR with primers designed to flank the IRE1-mediated splicing site (hamXBP1.19S: GGCCTTGTAATTGAGAACCAGGAG and mXBP1.14AS:

GAATGCCCAAAAGGATATCAGACTC). The amplification was carried out using the Q5® High-Fidelity 2X Master Mix (New England Biolabs), following the protocol described by Tung et al., (2024) (Tung *et al*, 2024). The PCR products corresponding to XBP1u (255 bp) and XBP1s (229 bp) were separated on a 3% agarose gel and visualised using SYBR Safe DNA stain (Invitrogen). Additionally, a hybrid amplicon of approximately 280 bp was detected. The proportion of XBP1s was quantified by measuring the intensity of gel bands using Fiji software (version 1.53c), with the hybrid band assumed to be equally composed of XBP1u and XBP1s.

### Generation of knockout cells using CRISPR-Cas9

Single guide RNAs (sgRNAs) targeting exon regions of *Slc33a1*, *Nat8*, and *Nat8b* in *Cricetulus griseus* were designed using CCTop (https://cctop.cos.uni-heidelberg.de:8043). These sgRNAs were subsequently cloned into one of the following CRISPR-Cas9 plasmid vectors: pSpCas9(BB)-2A-mCherry_V2 (UK1610) or pSpCas9(BB)-2A-mTurquoise (UK2915) as described by Ran et al. (Ran *et al*, 2013). Plasmid constructs were validated by DNA sanger sequencing to ensure correct insertion. For cell editing, previously described IRE1/ATF6α dual UPR reporter cells (Tung *et al*, 2024), IRE1/PERK dual UPR reporter cells (Sekine *et al*, 2015), 3×FLAG-mGL-ATF6α knock-in cells (Tung *et al*, 2024) and GFP-ATF6α_LD^S1P,S2Pmut^ stable cells (Tung *et al*, 2024) were transfected with 1 µg of the sgRNA/Cas9 plasmid using Lipofectamine LTX (Thermo Fisher Scientific) in a 6-well plate. After 48 h, cells with high mTurquoise or mCherry (depending on the reporter cells used to generate the knockout) fluorescence were sorted into 96-well plates at one cell per well using a MoFlo Cell Sorter (Beckman Coulter). Successful knockout clones were verified via Sanger sequencing (Supplemental Fig. S1B).

### Live cell confocal microscopy

Cells stably expressing GFP-ATF6α_LD^S1P,S2Pmut^ or cells having endogenous cgATF6α with an N-terminal 3×FLAG-mGL were cultured on 35 mm glass-bottom imaging dishes (MatTek) and pmScarlet_Giantin-C1 plasmid to label the Golgi apparatus in red. After 24 h of incubation post-transfection, cells were treated with either 4 mM 2-deoxy-D-glucose (2DG) for 3 h or 2 mM dithiothreitol (DTT) for 1 h, depending on the experimental condition. Live imaging was performed using a Zeiss LSM780 inverted confocal laser scanning microscope equipped with a temperature-and CO₂-controlled chamber (37°C, 5% CO₂). A ×64 oil immersion objective was used, with the pinhole set to 1 Airy unit. Fluorescent signals were captured using appropriate laser lines: 488 nm for GFP or mGL, 405 nm for DAPI, and 594 nm for mCherry. Co-localisation between different fluorescent markers within single cells was analysed using Volocity software (version 6.3, PerkinElmer) by calculating the Pearson correlation coefficient.

### Pulse-chase radiolabelling and Immunoprecipitation using GFP-Trap Agarose to detect Golgi modified glycans

Parental CHO-K1 cells and their *Slc33a1*-deleted derivatives were seeded in 12-well plates. After 18 h post-seeding, cells were transfected with the GFP-ATF6α_LD^S1P, S2Pmut^ construct (UK3116). Twenty-four hours post-transfection, cells were cultured in methionine/cysteine-deficient DMEM (GIBCO, Cat# 21013024) for 30 min to deplete endogenous amino acids. To initiate ATF6α trafficking from the ER to the Golgi, cells were treated with 300 nM Thapsigargin (Tg) for 30 min, with Tg treatment maintained throughout the subsequent radiolabelling step. Cells were then pulse-labelled for 10 min using 5.5 μCi/well of [³⁵S] methionine/cysteine (Expre35S Protein Labelling Mix), followed either by immediate harvesting or a chase period of up to 80 min. Post-pulse, media were collected, and cells were washed twice with ice-cold PBS containing 0.1 mg/ml cycloheximide to halt translation. Cells were then pelleted by centrifugation at 845 xg for 5 min at 4°C and lysed using Nonidet lysis buffer (150 mM NaCl, 50 mM Tris-HCl pH 7.5, 1% NP-40, 0.05 mM TCEP) supplemented with protease inhibitors (0.1 mg/ml cycloheximide, 1 mM PMSF, 4 µg/ml aprotinin, and 2 µg/ml pepstatin A). Following lysis, post-nuclear supernatants were obtained by centrifuging the lysates at 20,000 *xg* for 15 min at 4°C, and the soluble portion was collected. Radiolabelled proteins were subjected to digestion with Endo H, PNGase F, O-glycosidase, and neuraminidase individually or in combination. Equal volumes of the cleared digested lysates were incubated with 15-20 μl GFP-Trap Agarose (ChromoTek) preequilibrated in lysis buffer. The mixture was incubated for 2 h at 4°C with rotation. The beads were then recovered by centrifugation (845*xg*, 3 min) and washed four times with washing buffer (50 mM Tris-HCl, 1 mM EDTA, 0.1% Triton X-100). Proteins were eluted in 35 μl of 4×SDS-DTT sample for 10 min at 55°C, resolved on 12.5% SDS-PAGE gels, and visualized by autoradiography using a Typhoon biomolecular imager (GE Healthcare). Signal quantification was carried out using Fiji (ImageJ).

### Mass spectrometric analysis of ATF6α_LD^S1P,S2Pmut^ protein glycosylation

Parental and *Slc33a1*-deleted cells lines stably expressing GFP-ATF6α_LDS^1P,S2Pmut^ were cultured on four 140 mm plates. After 18 h post-seeding, cells were treated with 4 mM DTT for 1 h to induce ATF6 trafficking to the Golgi. Following treatment, cells were washed with ice-cold PBS, collected in PBS, centrifugated at 845 ×g for 5 min at 4°C and lysed in Nonidet lysis buffer supplemented with protease inhibitors, as previously described. Post-lysates supernatants were collected at 20,000 ×g for 15 min at 4°C. Equal volumes of the cleared lysates were incubated with 15-20 μl GFP-Trap Agarose (ChromoTek), pre-equilibrated in lysis buffer. The mixture was rotated for 2 h, at 4°C. The beads were then recovered by centrifugation (845 xg, 3 min) and washed four times with washing buffer (50 mM Tris-HCl, 1 mM EDTA, 0.1% Triton X-100). Beads were washed 3 times with PBS to remove the detergent traces, and then resuspended in 50 ul of PBS for further processing by mass spectrometry.

20 µL of beads was transferred to 150 µL of 10mM dithiothreitol in 50mM ammonium bicarbonate solution in Lo-Bind tubes (Eppendorf) and incubated at 60°C for 1 h. Vials were mixed every 15 min to resuspend the beads during the reduction process. Reduced cysteine bonds were capped by adding 40 µL of 100 mM iodoacetamide in 50 mM ammonium bicarbonate and incubated in the dark for 30 min. Vials were brought into the light for 20 min before 30 µL of 50 mM ammonium bicarbonate containing 1 µg of Trypsin Gold (Promega). Samples were incubated overnight at 37 °C and quenched with 20 µL of 1% formic acid in water (v/v) and transferred to LC-MS vials for analysis.

Digested samples were initially analysed on a Waters M-Class LC system connected to a SCIEX ZenoToF mass spectrometer. Sample (20 µL) was injected onto a Waters NanoEase 0.18 x 20 mm Symmetry C18 trap column and then switched in line with a Waters NanoEase 0.075 x 150 mm HSS T3 nano column. Peptides were eluted into the mass spectrometer using a gradient of 2 to 40% ACN over 51 min flowing at 0.5 µL/min. Peptides were analysed using an IDA based analysis with the top 50 selected for fragmentation, with an exclusion period of 12 seconds. LC-MS/MS files were searched using PEAKS 12.5 (BSI) against a custom database containing the ATF6α_LD^S1P,S2Pmut^ protein, using a fixed cysteine carbamidomethylation modification, and variable modifications of oxidised methionines and deamidated asparagines and glutamines. The PEAKS search was then run with a Glycan search, which contains 1828 basic glycans.

The extracts were reanalysed on a Waters M-Class LC system linked to a Waters TQ-XS triple quadrupole mass spectrometer with an IonKey interface. Peptides were injected onto a Waters NanoEase 0.3 x 50mm C18 trap column before switching in line with a 0.15 x 50mm HSS T3 iKey. Peptides were separated over a gradient of 2 to 55% ACN over 37 min flowing at 1 µL/min. The triple quadrupole monitored for precursor ions of 2 glycosylated peptide variants and monitored characteristic oxonium ions relating to glycosylated peptides as well as the HVVEFGGENLYFQSAK (HVVE) backbone peptide.

| Peptide | Q1 | Q3 | Charge | CE |
| --- | --- | --- | --- | --- |
| HVVE backbone | 608.97 | 580.31, 743.37*, 855.40, 970.45 | 3+ | 25 |
| HEX8 peptide | 1022.12 | 163.06, 204.08, 366.14* | 3+ | 40 |
| NeuAc2 peptide | 928.76 | 204.08, 274.09*, 366.14 | 4+ | 30 |

This table shows precursor (Q1) and product ion (Q3) information and collision energy (CE) for two glycopeptides and the HVVE backbone peptide. The product ion with an asterisk was used to generate quantitative data. Other product ions were monitored for qualitative data only.

### Key Resources Tables

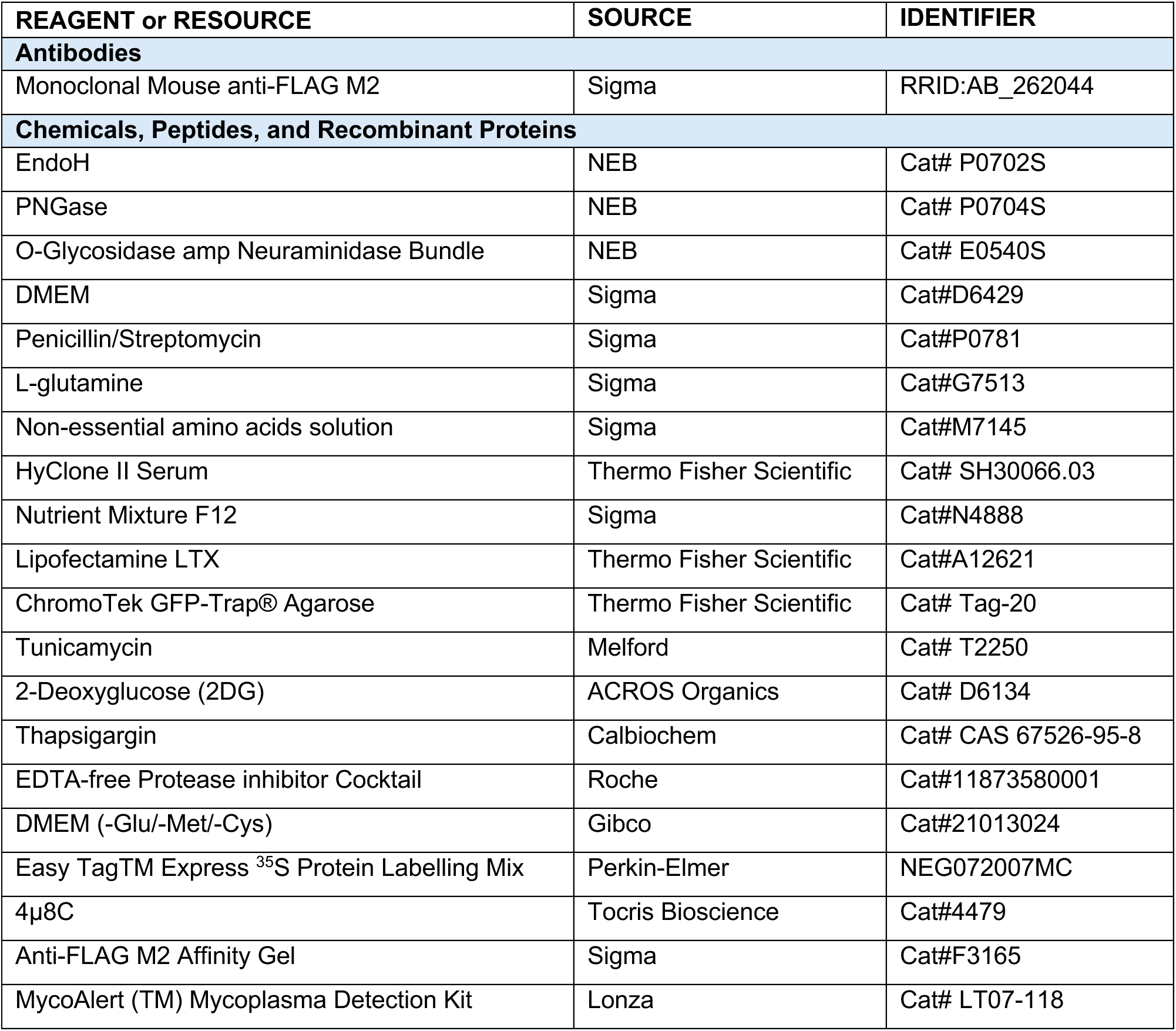
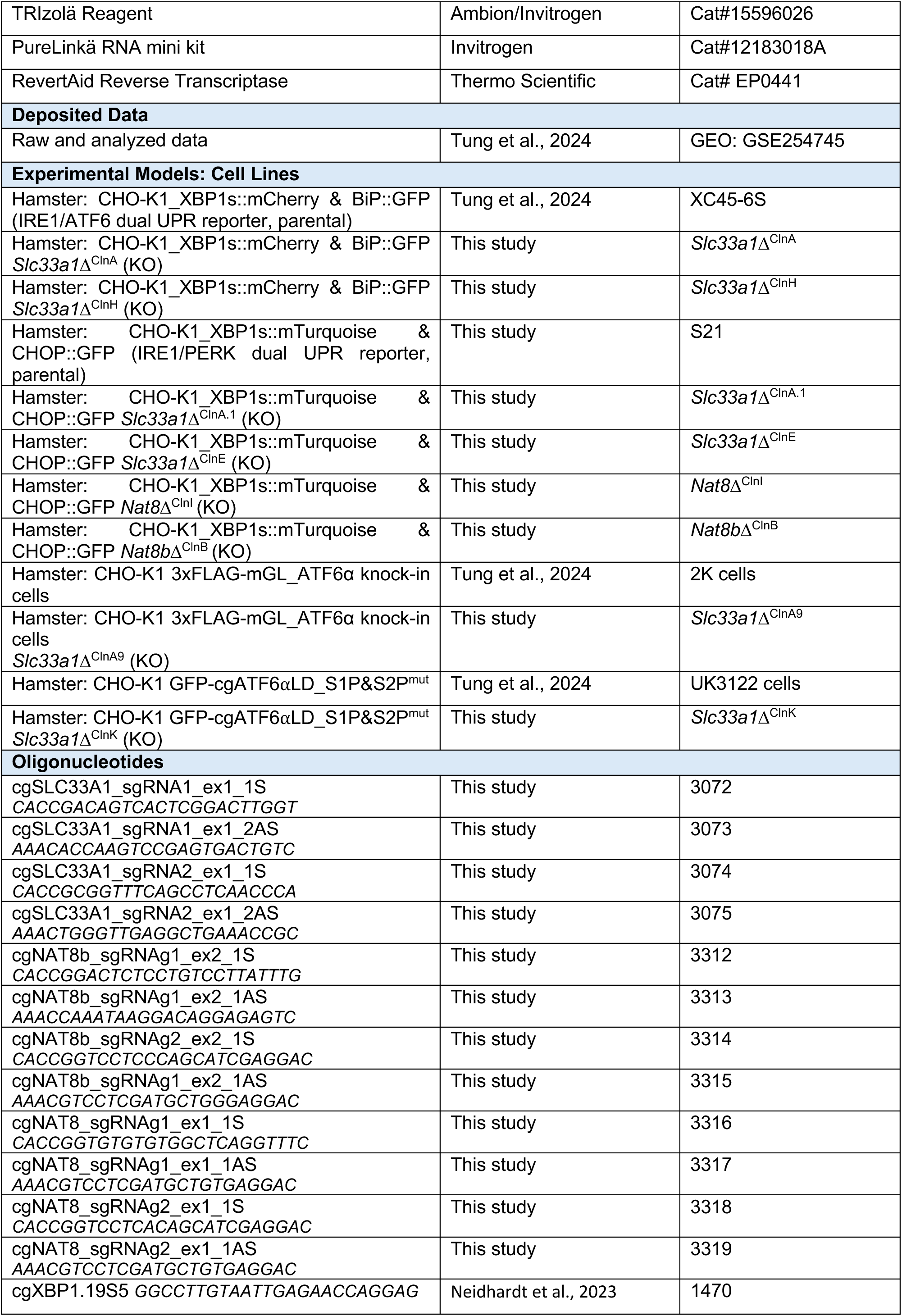
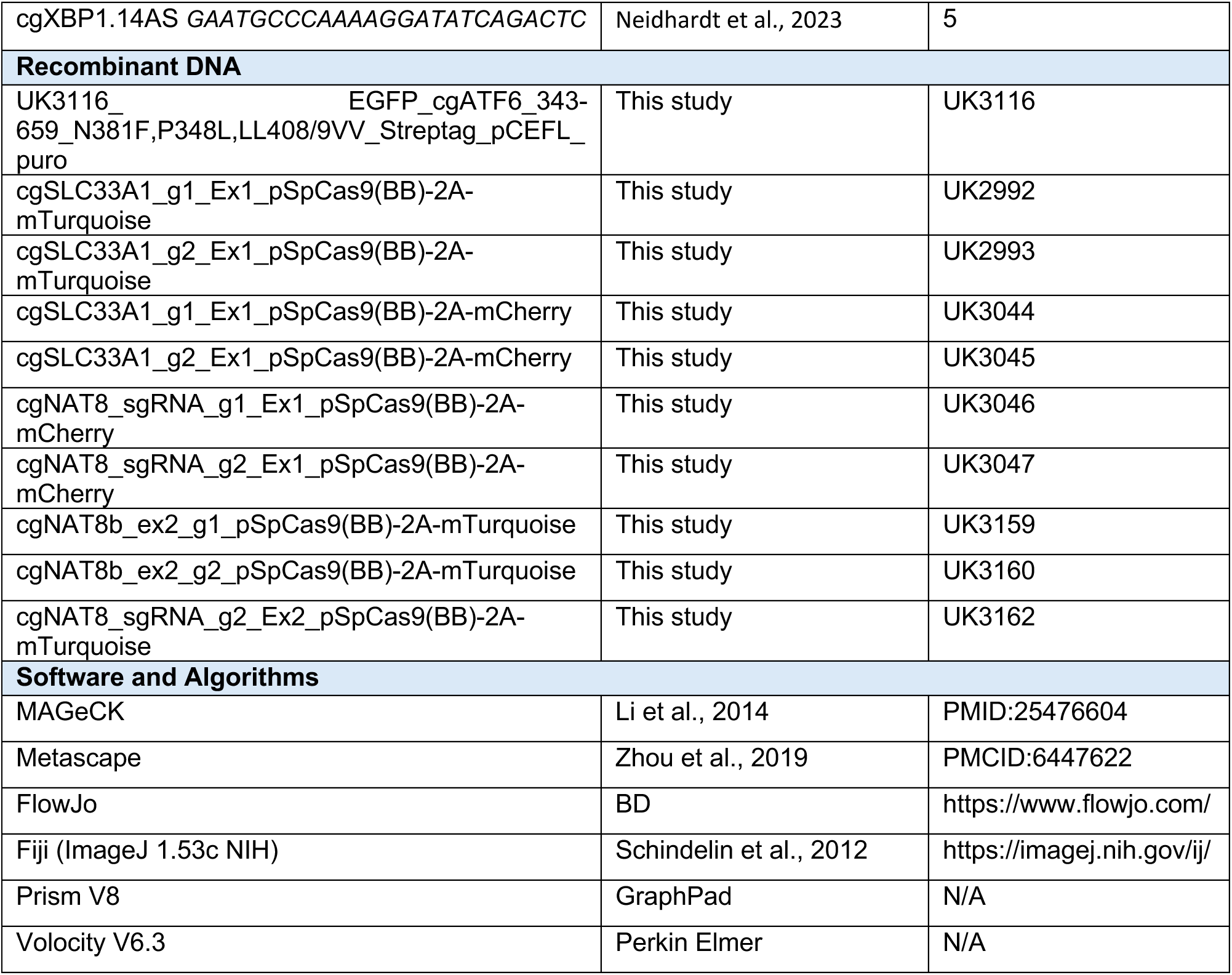

## Author contributions

**Ginto George**: Conceptualisation; Data curation; Formal analysis; Validation; Investigation; Methodology; Writing-original draft; Writing-review and editing. **Heather P Harding**: Investigation; Methodology; Writing-review and editing. **Richard Kay:** Methodology**. David Ron**: Conceptualisation; Resources; Supervision; Funding acquisition; Visualization; Writing-original draft; Writing-review and editing. **Adriana Ordoñez**: Conceptualisation; Data curation; Formal analysis; Methodology; Supervision; Funding acquisition; Writing-original draft; Writing-review and editing.

## Supporting information

Supplemental Figure S1. CRISPR-Cas9-mediated targeting and validation of Slc33a1 disruption in CHO-K1 cells. Related to Figure 1.

Supplemental Figure S2. Specific activation of IRE1 signalling with reduced ATF6a activation in Slc33a1-deleted clones. Related to Figure 2.

Supplementary Figure S3. Constitutive Golgi localisation of ATF6a in Slc33a1-deleted cells. Related to Figure 4.

Supplementary Figure S4. Analysis of Golgi-modified N-glycans of ATF6 by mass spectrometry. Related to Figure 5.

## Acknowledgements

We thank the CIMR flow cytometry core facility team (Reiner Schulte and Gabriela Grondys-Kotarba) and the microscopy team (Matthew Gratian and Mark Bowen) for technical support. Tianyu Wen and Stuart Haslam (Imperial College London) for assistance in glycobiology in early phases of this research. Supported by a Wellcome Trust Principal Research Fellowship to DR (Wellcome 224407/Z/21Z) and by the Spanish Ministry of Science, Innovation and Universities to AO (RYC2022-035365-I).

## Supplemental Figures

**Supplemental Figure S1.**
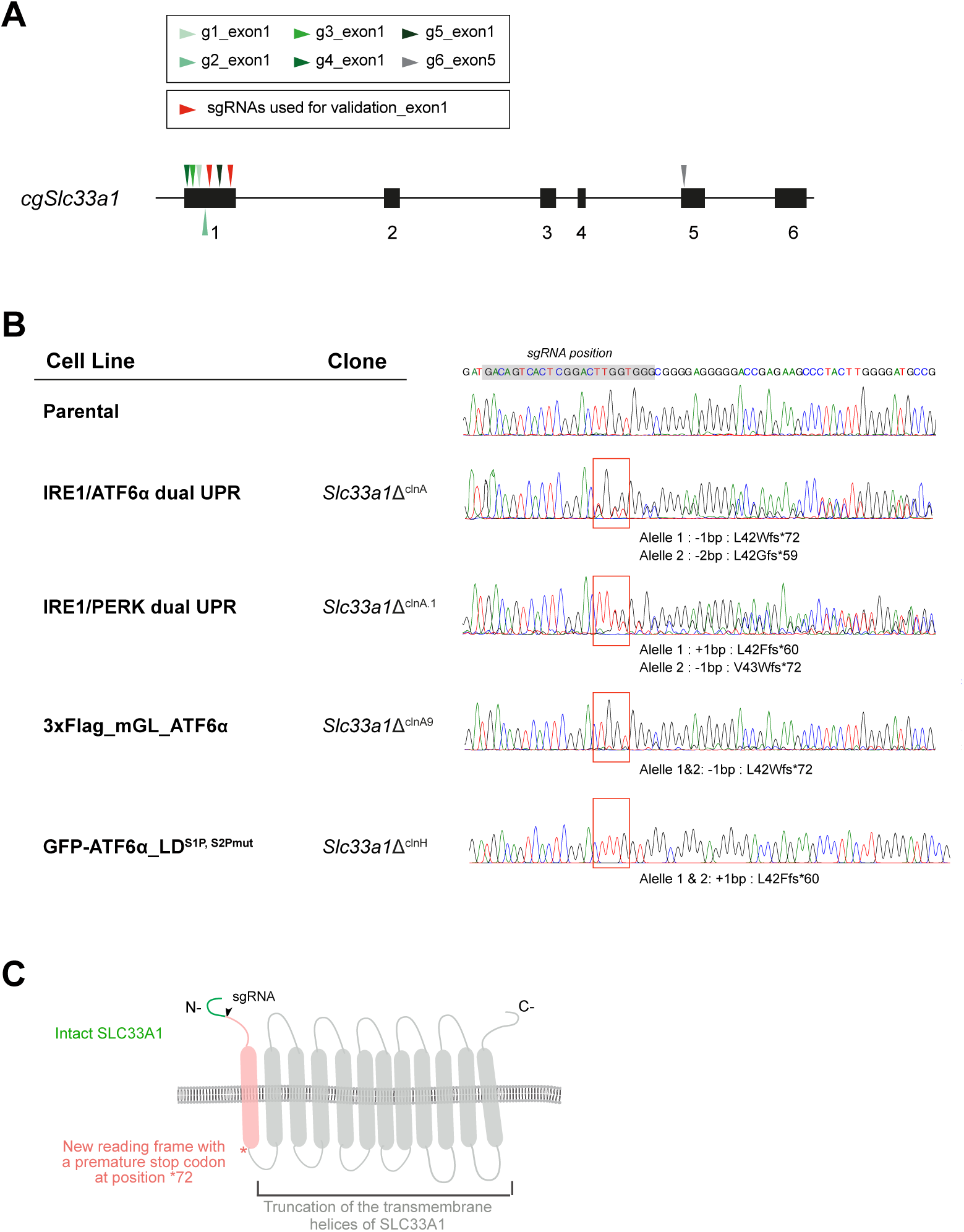
***CRISPR-Cas9-mediated targeting and validation of Slc33a1 disruption in* CHO-K1 cells*. Related to* *Figure 1*. (A)** Schema of the *Slc33a1* locus in *Cricetulus griseus* (*cg*), indicating the positions of the six sgRNAs (g1-g6) included in the CRISPR-Cas9 library (arrowheads in green and grey). Red arrowheads denote the two sgRNAs selected for validation. **(B)** Sanger sequencing chromatograms displaying the targeted region of exon 1 of the *cgSlc33a1* locus. These confirm the generation of early premature stop codons in all *Slc33a1*-deleted cells generated in this study. The grey shading indicates the sgRNA target sequence within *Slc33a1* locus, while the red box indicates the mutation start point. **(C)** Cartoon representation of the SLC33A1 structure, indicating the intact portion of the protein in the *Slc33a1*-deleted cells (green), the mutation start point (black arrow), the resulting new reading frame (in red), and the position of the stop codon beyond which the remainder of the protein is truncated (grey).

**Supplemental Figure S2.**
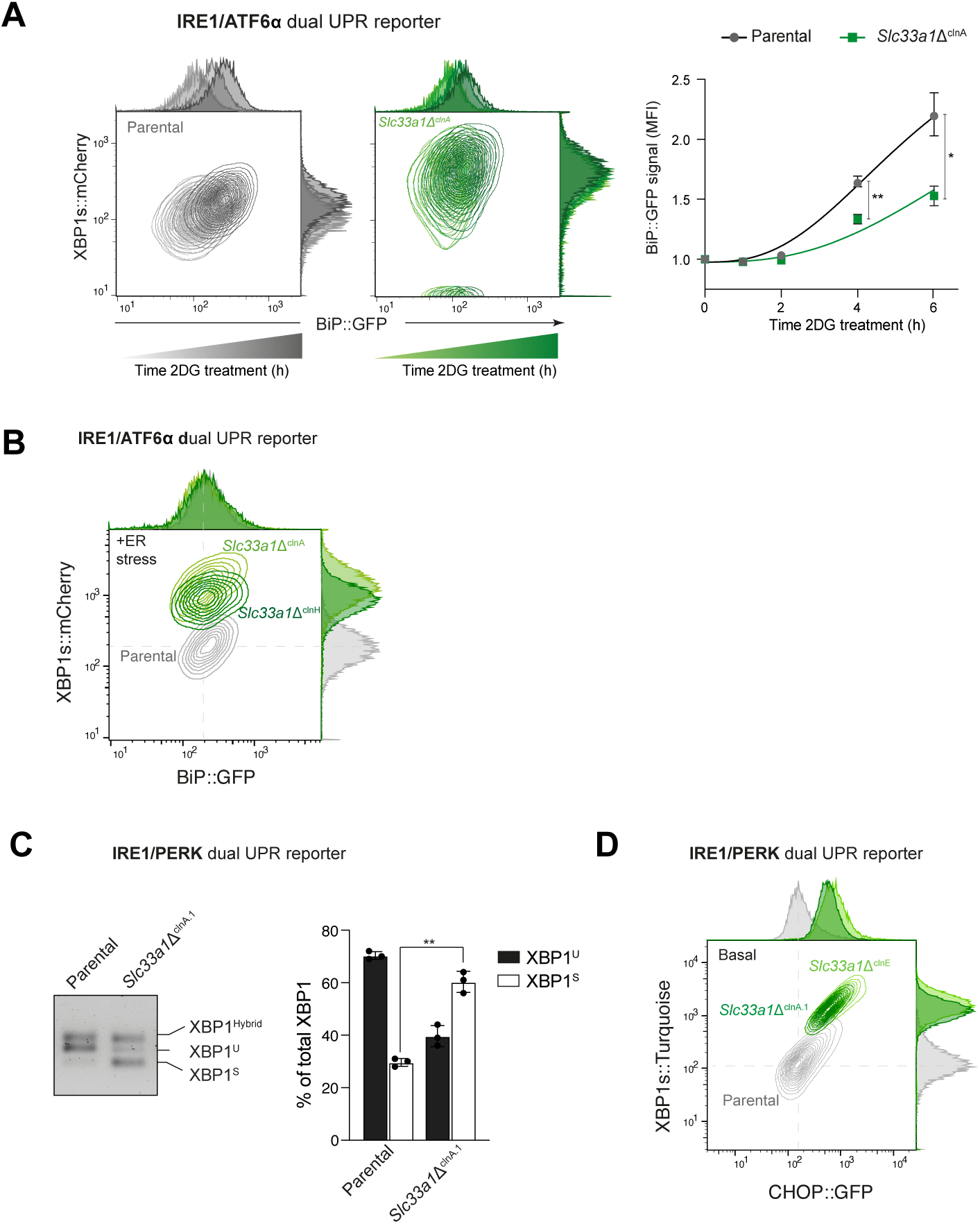
***Specific activation of IRE1 signalling with reduced ATF6α activation in Slc33a1Δ clones. Related to* *Figure 2*. (A)** Related to Figure 2A. Two-dimensional contour plots depicting the XBP1s::mCherry and BiP::GFP signals in parental IRE1/ATF6α dual UPR reporter cells (left panel, grey gradient) and in the *Slc33a1*Δ^clnA^ derivative clone (right panel, green gradient) treated with 2-deoxy-D-glucose (2DG, 4 mM) over a time course up to 6 h. The normalised fold induction of the median BiP::GFP signal intensity over the time course for each genotype is shown in a graph as mean ± SD from three independent experiments. *p < 0.05 and **p < 0.01, according to 2way ANOVA test followed by a Bonferroni *post hoc* test. **(B)** Related to Figure 2A. Two-dimensional contour plots depicting XBP1s::mCherry and BiP::GFP signals in parental IRE1/ATF6α dual UPR reporter cells (grey) and two derivative *Slc33a1*-deleted clones: *Slc33a1*Δ^clnA^ (light green; previously shown in Fig. 2A) and an additional *Slc33a1*Δ^clnH^ (dark green). Data shown reporter intensity under ER stress condition with Tunicamycin (Tm, 2.5 µg/ml, 6 h). Data are representative from two independent experiment. **(C)** Related to Figure 2D. <u>Left panel:</u> Representative agarose gel showing XBP1 cDNA isoforms detected via RT-PCR in IRE1/PERK dual UPR reporter parental cells and *Slc33a1*Δ^clnA.1^ cells under basal conditions. The migration of the unspliced (XBP1^U^), spliced XBP1 (XBP1^S^) and hybrid (XBP1^Hybrid^, containing one strand of XBP1^U^ and one strand of XBP1^S^) stained DNA fragments is indicated. <u>Right panel:</u> Plot of the percentage of XBP1^U^ and XBP1^S^ fractions of XBP1 mRNA in the two genotypes. Bars represent mean ± SD, and the values of the three replicates as black dots (**p < 0.01; two-sided unpaired Welch’s t-test). **(D)** Related to Figure 2D. Two-dimensional contour plots depicting XBP1s::mCherry and CHOP::GFP signals in parental IRE1/PERK dual UPR reporter cells (grey) and two derivative *Slc33a1*-deleted clones: *Slc33a1*Δ^clnA.1^ (dark green; previously shown in Fig. 2D) and an additional *Slc33Aa1*Δ^clnE^ (light green). Data shown reporter intensity under basal conditions. Data are representative from three independents experiments.

**Supplementary Figure S3.**
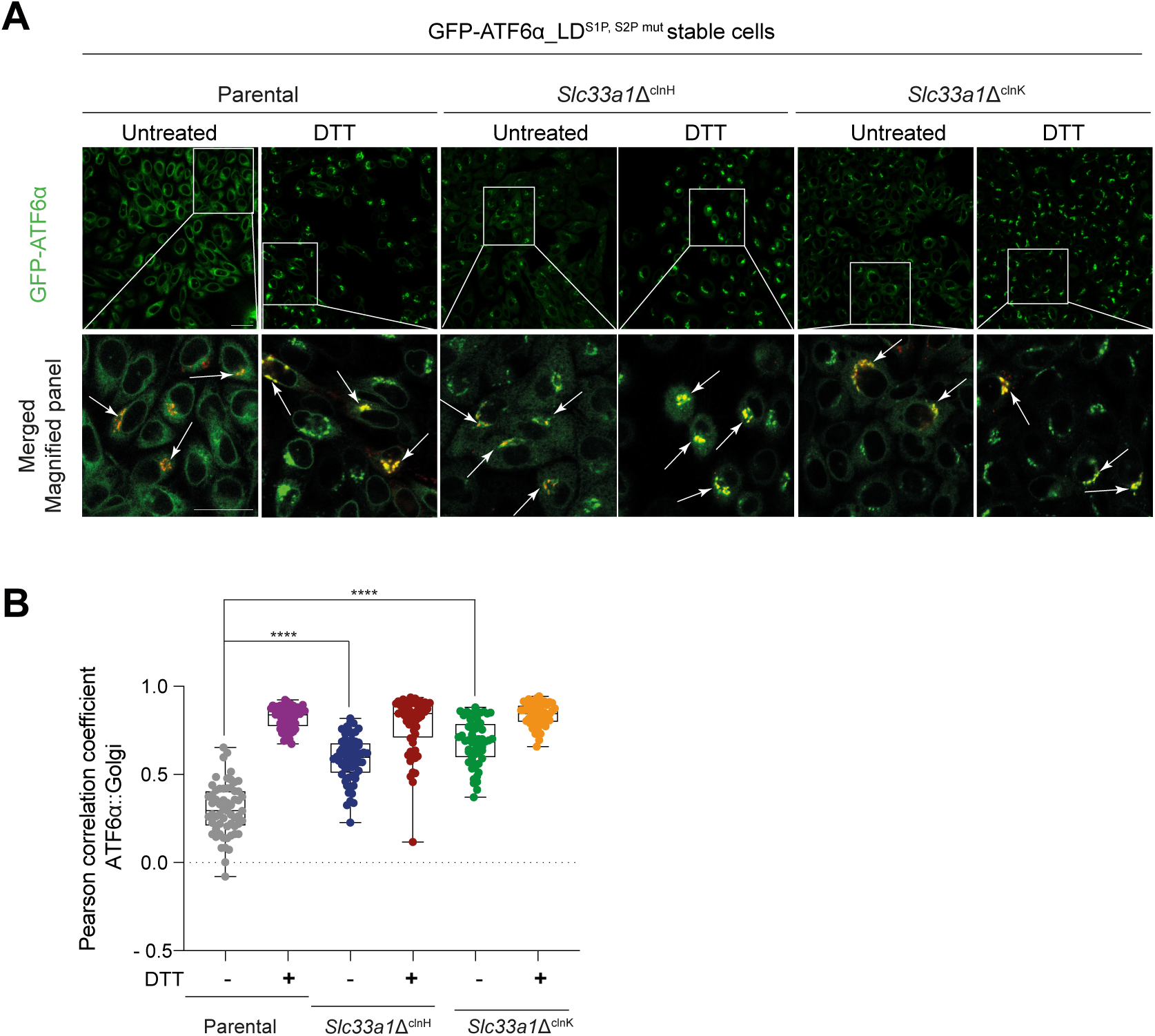
***Constitutive Golgi localisation of ATF6α in Slc33a1Δ Cells. Related to* *Figure 4*. (A)** Related to Figure 4D-F. Representative live-cell confocal microscopy images of CHO-K1 stably expressing GFP-ATF6α_LD^S1P,S2Pmut^ parental cells and two derivative *Slc33a1*-deleted clones: *Slc33a1*Δ^clnH^ (previously shown in Fig. 4D-F) and an additional *Slc33a1*Δ^clnK^. mGL-ATF6α is shown in green. Cells were transiently transfected with the pmScarlet_Giantin-C1 plasmid to label the Golgi apparatus in red (Merge panel). To promote ATF6α localisation to the Golgi, cells were treated with DTT (2 mM, 1 hr). Insets show magnified regions, with arrows highlighting co-localisation of GFP-ATF6α_LD^S1P,S2Pmut^ with Giantin. Scale bar: 20 μm. **(B)** Pearson’s correlation coefficients were used to quantify co-localisation of GFP-ATF6α_LD^S1P,S2Pmut^ with Giantin in parental cells (n > 50) and both *Slc33a1*-deleted clones (n > 50). Data are presented as a box-and-whisker plot, including all individual values along with minimum and maximum intensities. Co-localisation analysis was performed using Volocity software, and statistical significance was determined using a two-sided unpaired Welch’s t-test (****p < 0.0001).

**Supplementary Figure S4.**
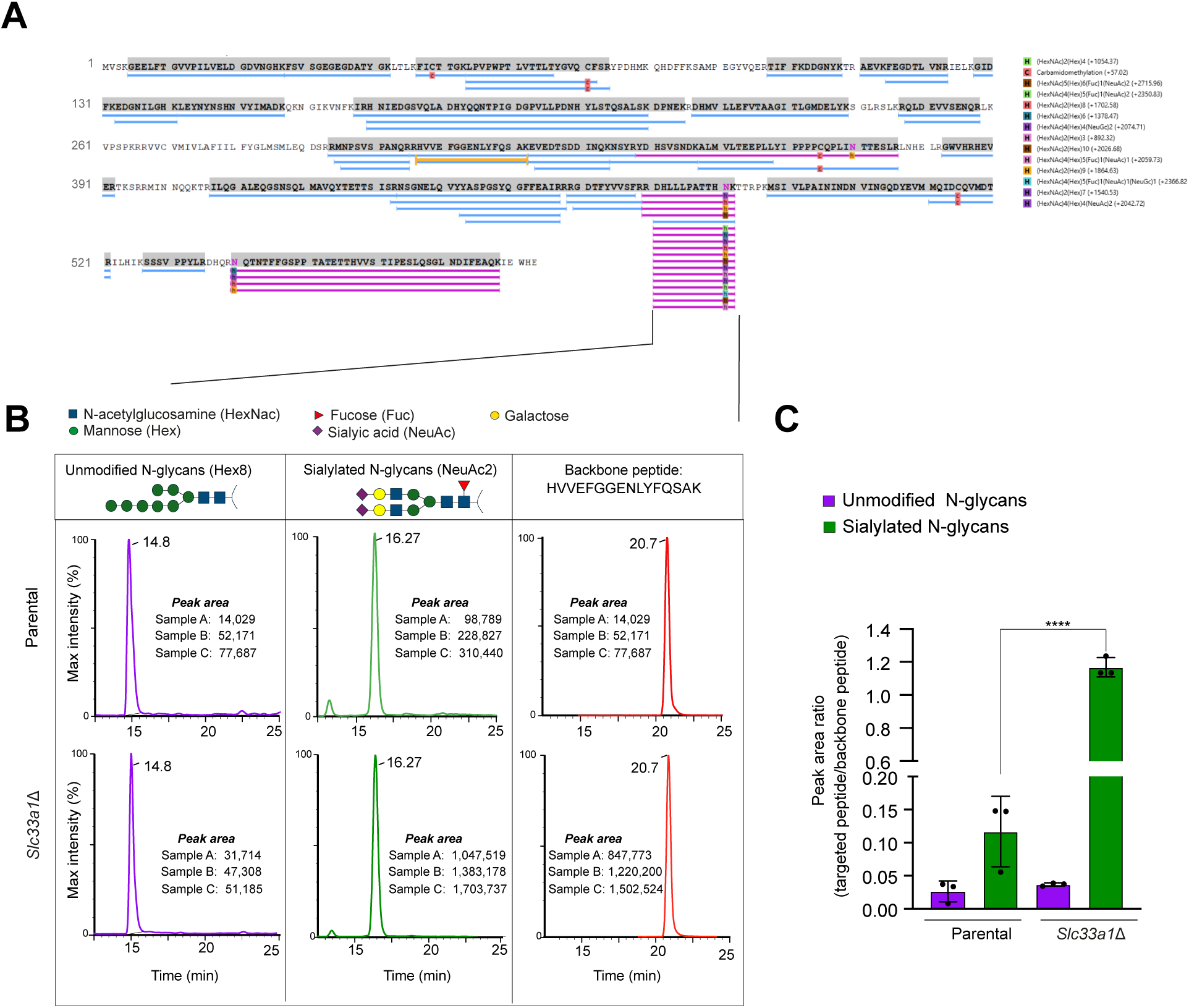
Analysis of Golgi-modified N-glycans of ATF6 by mass spectrometry. Related to. **Figure 5. (A)** Protein coverage map displaying peptide identifications of the selected protein (GFP-ATF6LD) mapped onto the protein sequence. Each blue bar indicates an identified peptide sequence. Peptides carrying N-glycosylation modifications are highlighted in pink, while the peptide used as a backbone reference is shown in yellow. **(B)** Extracted ion chromatograms from a typical sample analysed on the Waters TQ-XS, showing the total ion current for the HVVE backbone peptide, the unmodified N-glycan peptide (denoted as HEX8) and the sialylated N-Glycan peptide (denoted as NeuAc2) eluting at 20.7, 14.8 and 16.27 min respectively. Next to each chromatogram, the peak areas for three replicates (A–C) of the corresponding peptide are shown for both genotypes: parental and *Slc33a1*Δ. **(C)** Quantification of the targeted peptide expresses as the peak area ratio of the glycosylated peptide to that of the backbone peptide. Bars represent mean ± SD, with individual replicate values shown as black dots (****p < 0.0001; two-sided unpaired Welch’s t-test).

