## Supplementary figures and images for "Metabolite import via SLC33A1 enables ATF6 activation by endoplasmic reticulum stress"

### Supplemental Figure S1. CRISPR-Cas9-mediated targeting and validation of Slc33a1 disruption in CHO-K1 cells. Related to Figure 1.

#### Supplemental Fig. S1. Related to Fig. 1

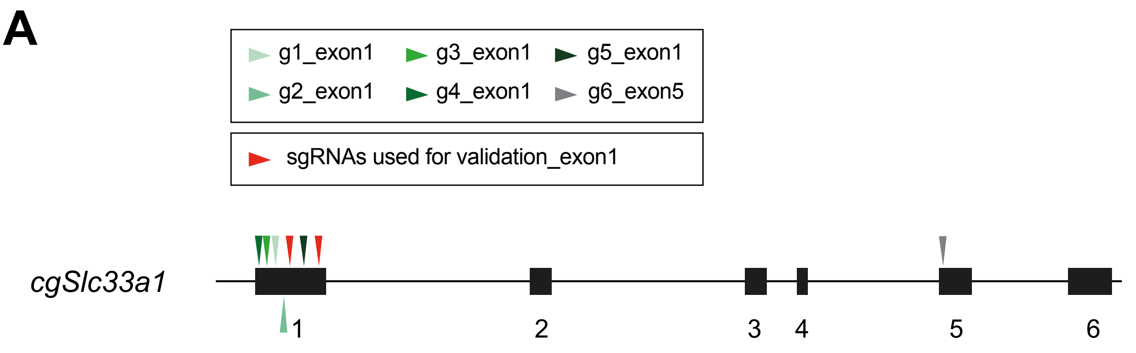


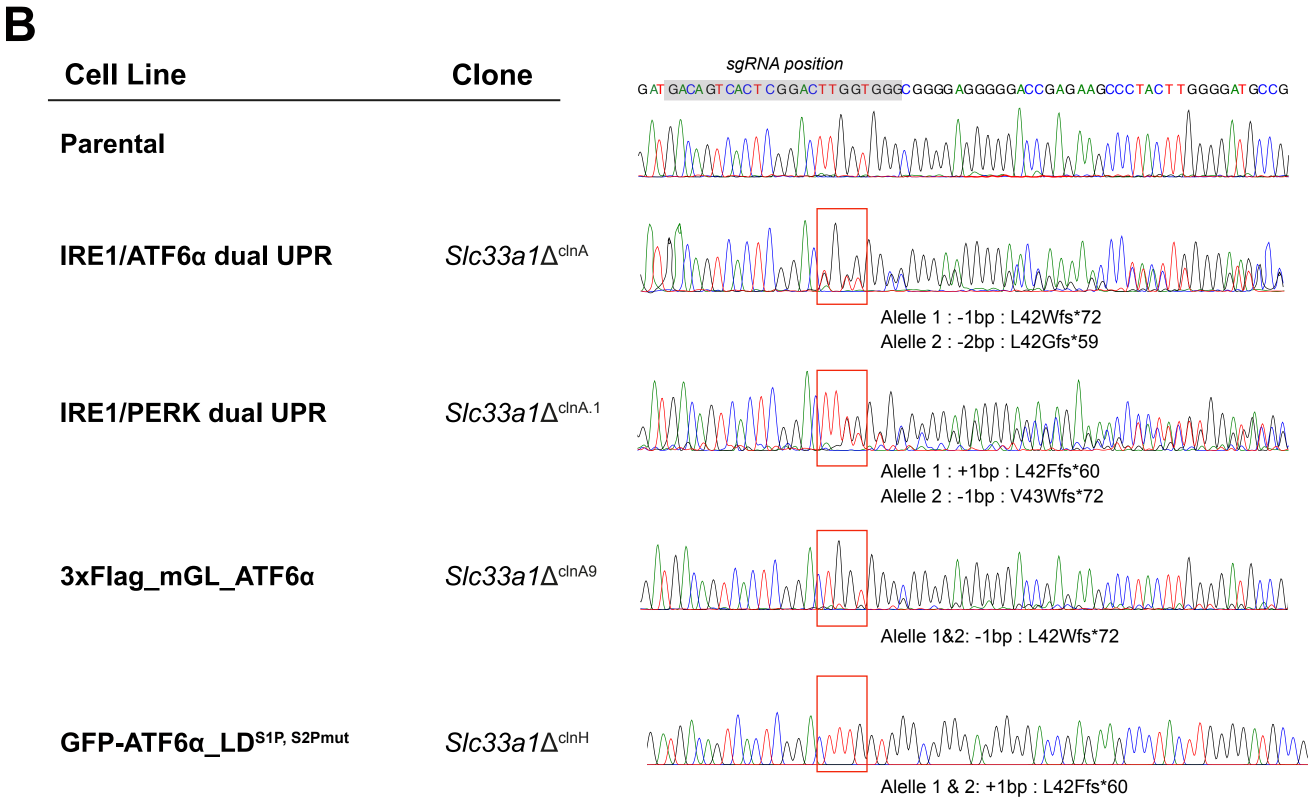


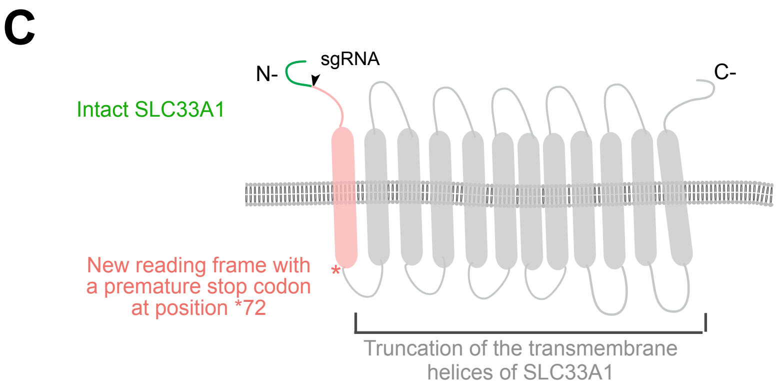

### Supplemental Figure S2. Specific activation of IRE1 signalling with reduced ATF6a activation in Slc33a1-deleted clones. Related to Figure 2.

#### Supplemental Fig. S2. Related to Fig. 2

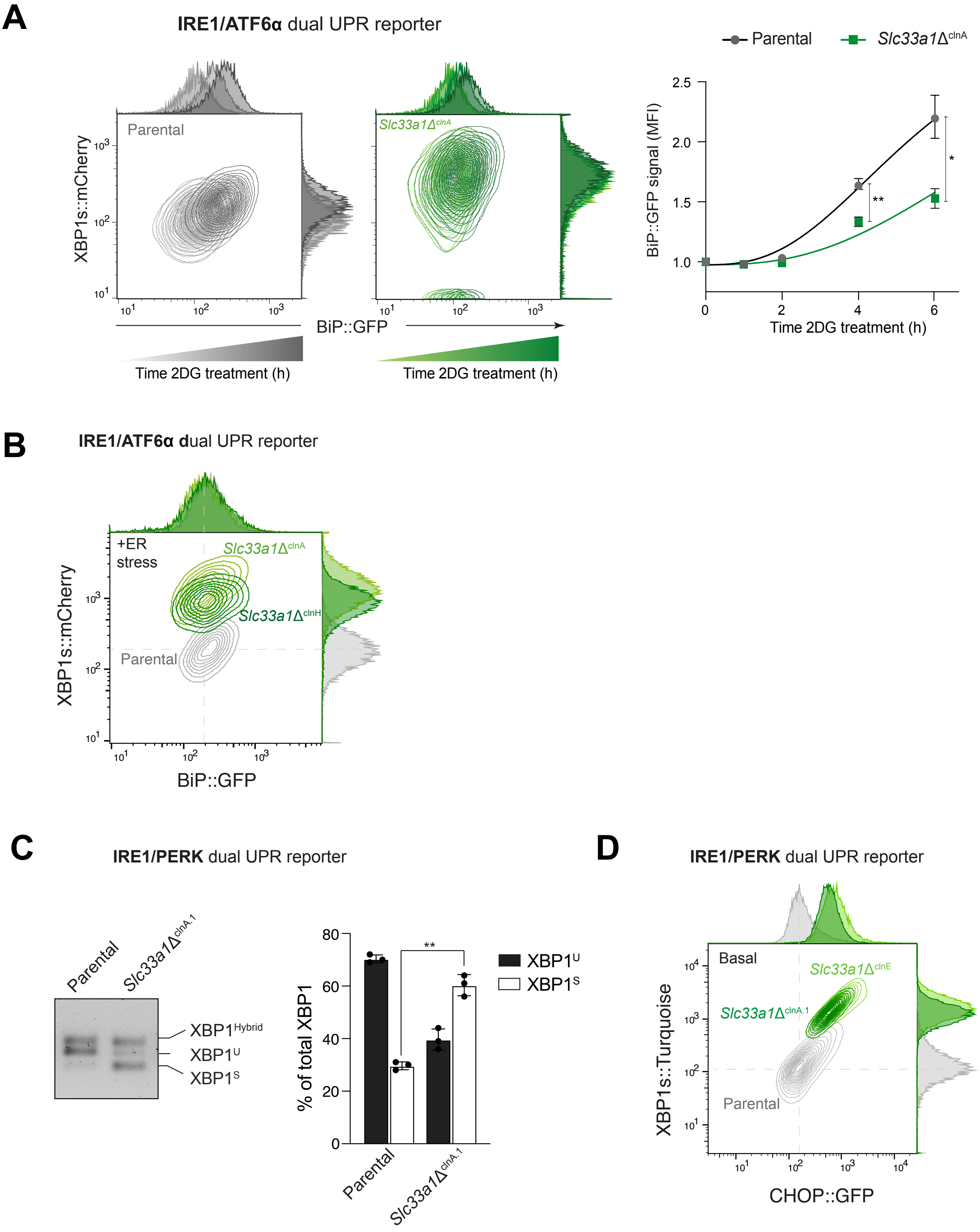

### Supplementary Figure S3. Constitutive Golgi localisation of ATF6a in Slc33a1-deleted cells. Related to Figure 4.

#### Supplemental Figure S3. Related to Fig. 4

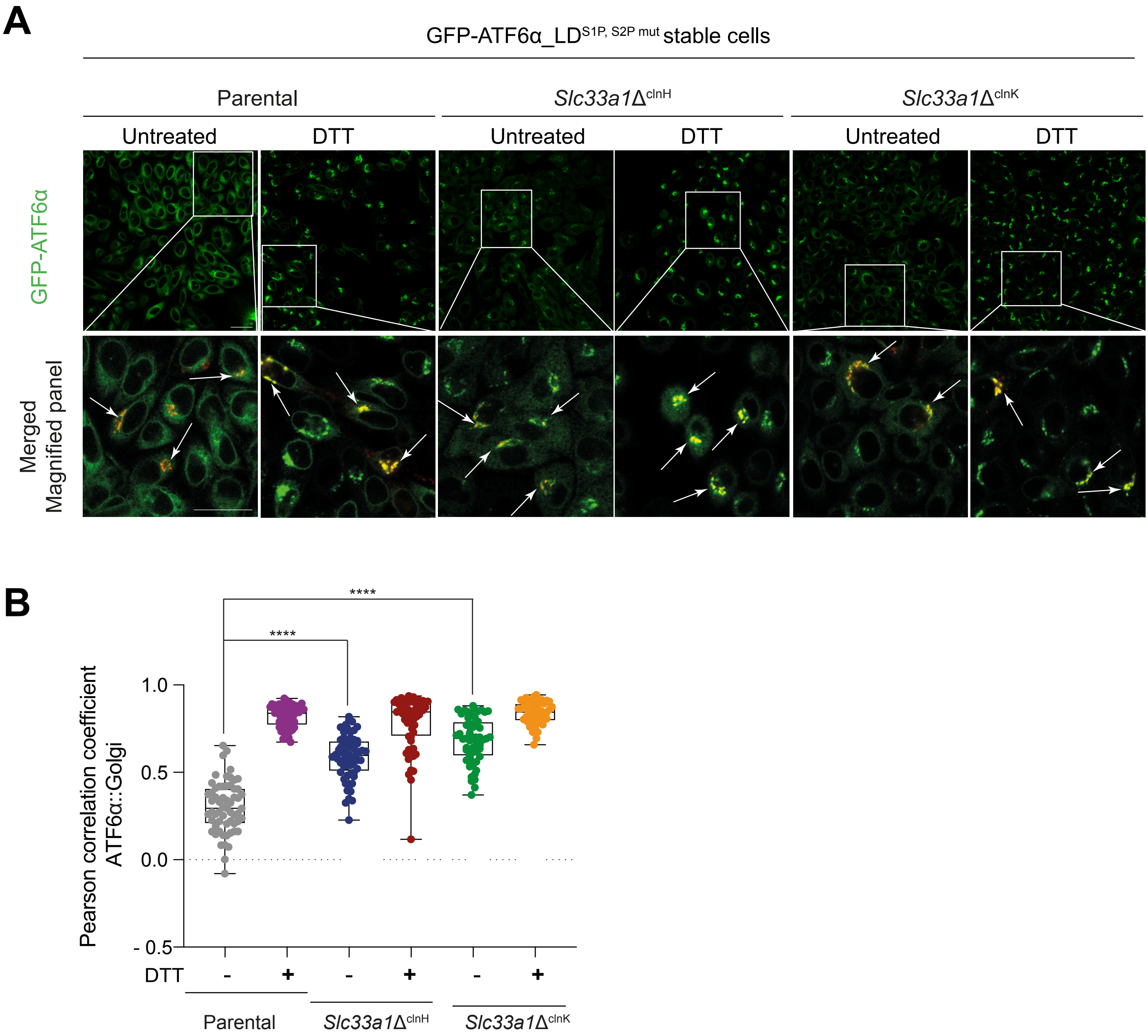

### Supplementary Figure S4. Analysis of Golgi-modified N-glycans of ATF6 by mass spectrometry. Related to Figure 5.

#### Supplemental Figure S4. Related to Fig. 5

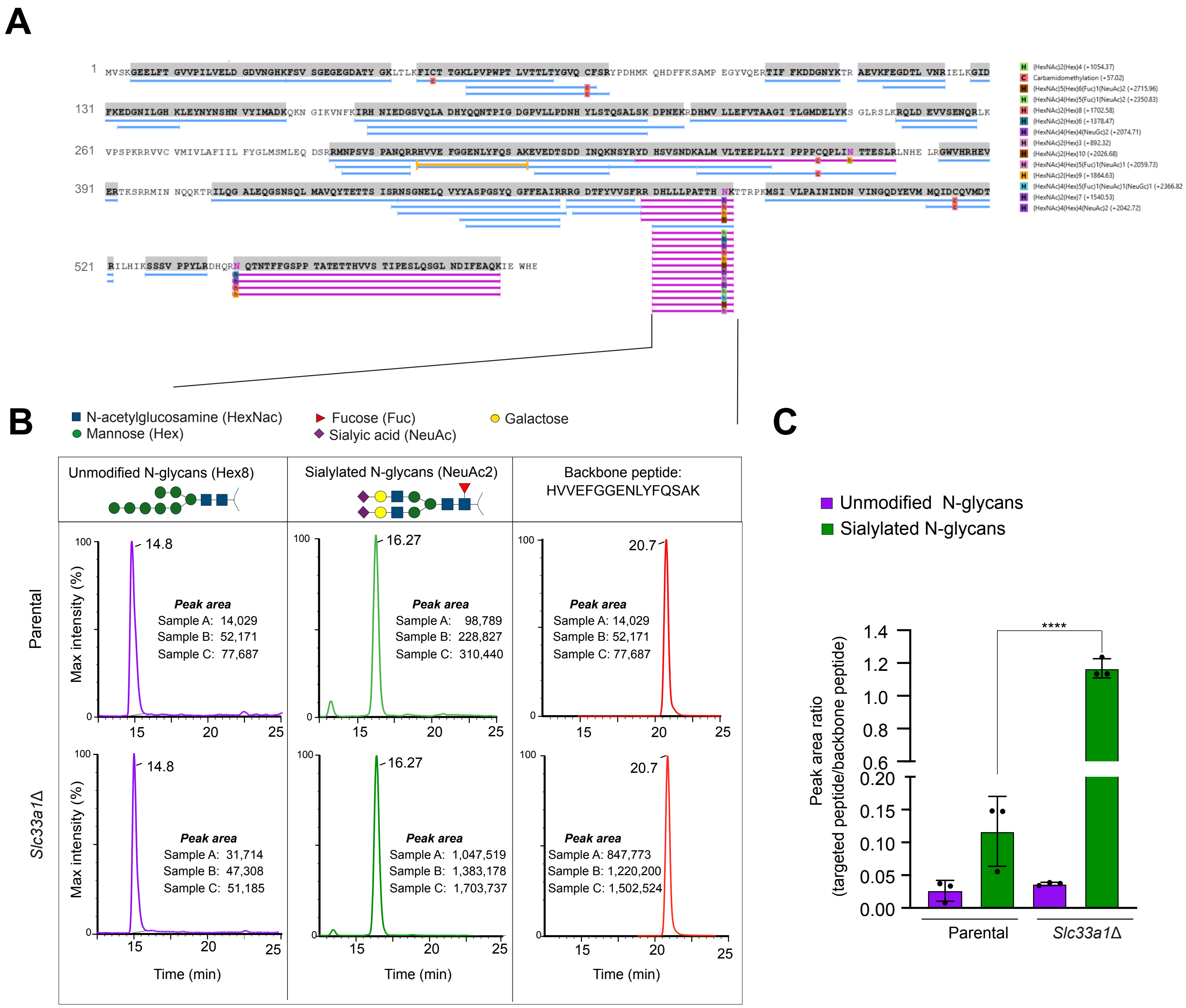
